# Democratizing biomedical simulation through automated model discovery and a universal material subroutine

**DOI:** 10.1101/2023.12.06.570487

**Authors:** Mathias Peirlinck, Kevin Linka, Juan A. Hurtado, Gerhard A. Holzapfel, Ellen Kuhl

**Affiliations:** Department of Biomechanical Engineering, Delft University of Technology · Delft · the Netherlands; Institute for Continuum and Material Mechanics, Hamburg University of Technology · Hamburg · Germany; Dassault Systèmes, ovidence · Rhode Island · United States; Institute of Biomechanics, Technical University of Graz · Graz · Austria; Norwegian University of Science and Technology · Trondheim Norway; Department of Mechanical Engineering Stanford University · Stanford · California · United States

**Keywords:** constitutive neural networks, machine learning, automated model discovery, finite element analysis, hyperelasticity, cardiovascular mechanics

## Abstract

Personalized computational simulations have emerged as a vital tool to understand the biomechanical factors of a disease, predict disease progression, and design personalized intervention. Material modeling is critical for realistic biomedical simulations, and poor model selection can have life-threatening consequences for the patient. However, selecting the best model requires a profound domain knowledge and is limited to a few highly specialized experts in the field. Here we explore the feasibility of eliminating user involvement and automate the process of material modeling in finite element analyses. We leverage recent developments in constitutive neural networks, machine learning, and artificial intelligence to discover the best constitutive model from thousands of possible combinations of a few functional building blocks. We integrate all discoverable models into the finite element workflow by creating a universal material subroutine that contains more than 60,000 models, made up of 16 individual terms. We prototype this workflow using biaxial extension tests from healthy human arteries as input and stress and stretch profiles across the human aortic arch as output. Our results suggest that constitutive neural networks can robustly discover various flavors of arterial models from data, feed these models directly into a finite element simulation, and predict stress and strain profiles that compare favorably to the classical Holzapfel model. Replacing dozens of individual material subroutines by a single universal material subroutine–populated directly via automated model discovery–will make finite element simulations more user-friendly, more robust, and less vulnerable to human error. Democratizing finite element simulation by automating model selection could induce a paradigm shift in physics-based modeling, broaden access to simulation technologies, and empower individuals with varying levels of expertise and diverse backgrounds to actively participate in scientific discovery and push the boundaries of biomedical simulation.

## 1 Motivation

Computational simulations play a pivotal role in understanding and predicting the biomechanical factors of a wide variety of cardiovascular diseases [8, 60, 61]. In vascular medicine, knowing the precise stress and strain fields across the vascular wall is critical for understanding the formation, growth, and rupture of aneurysms [28]; for identifying high-risk regions of plaque formation, rupture, and thrombosis [48]; and for optimizing stent materials, structure, and deployment in aortic stenosis [31]. The accurate simulation of cardiovascular disease is a complex challenge that requires collective efforts across a multitude of disciplines including cardiovascular medicine, applied mathematics, biomechanics, and computer science [44]. Clearly, it is impossible that everyone has a specialized training in material modeling and an in-depth knowledge in finite element simulation [10]. However, selecting a poor material model does not only jeopardize the success of the entire simulation, but can have life-threatening consequences for the patient. *The objective of this manuscript is to explore whether and how we can automate the process of material modeling and its integration into a finite element analysis*.

### Constitutive neural networks autonomously discover material models from data

Throughout the past couple of years, two alternative strategies have emerged to discover models directly from data: non-interpretable and interpretable approaches. *Non-interpretable approaches* closely follow traditional neural networks and typically discover functions of rectified linear unit, softplus, or hyperbolic tangent type [19]. Representatives of this category are tensor basis Gaussian process regression [14, 15], plain constitutive artificial neural networks [26, 32], and neural ordinary differential equations [54,56]. These approaches are straightforward to implement, provide an excellent approximation of the data, and can be integrated manually within finite element software packages [19,55]. However, the models and parameters that these methods learn are non-interpretable, meaning they provide little insight into the underlying material behavior [46]. *Interpretable approaches* discover models that are made up of a library of functional building blocks that resemble traditional constitutive models. Representatives of this category are sparse regression [11, 12], symbolic regression [2], and custom-designed constitutive neural networks [34, 53], the method we adopt here. These approaches a priori satisfy material objectivity, material symmetry, thermodynamic consistency, and polyconvexity [30], and autonomously discover free energy functions that feature popular constitutive terms and parameters with a clear physical interpretation. By design, all three translate smoothly into user material subroutines for a finite element analysis [1], and we could adopt any of these interpretable approaches. Here, for illustrative purposes, we use a custom-designed constitutive neural network to discover the best constitutive model for aortic tissue from thousands of possible combinations of a few functional building blocks [33]. We integrate *all* discoverable models into the finite element workflow by creating a universal material subroutine that contains 2^16^ = 65,536 constitutive models, made up of 16 individual terms [35]. We train and test our network with biaxial extension tests of the medial and adventitial layers of a human aorta, and discover various flavors of arterial models from the experimental data [25, 40].

### Model discovery is a non-convex optimization problem with multiple local minima

Unfortunately, in practice, the sixteen terms of the network tend to span a parameter space with multiple local minima, the network often discovers non-sparse solutions, and model discovery can become non-unique [38]. A successful strategy to address these limitations is *L*_*p*_ regularization [11], a powerful method to shrink the parameter space by penalizing the loss function with a penalty term that consists of the *L*_*p*_ norm of the parameter vector, weighted by a penalty parameter [5]. To illustrate the potential of *L*_*p*_ regularization, we first use *L*_0_ regularization, or discrete combinatorics [13], to discover the best-in-class one- and two-term models and then use *L*_1_ regularization, or lasso [58], to systematically reduce the number of terms. This allows us to discover a suite of different models for the media and for the adventitia, and learn about their structural and mechanical differences [21].

### Mechanical differences in media and adventitia modulate the pathogenesis of cardiovascular disease

Understanding the subtle structural and mechanical distinctions between the media and adventitia layers of the aorta is crucial for comprehending vascular health and disease [28]. The media is rich in smooth muscle cells and elastin fibers to provide elasticity and contractility, and facilitate hemodynamic function, while the adventitia is made up primarily of fibroblasts and collagen fibers to provide structural support [23]. Disruptions in the delicate structural and mechanical balance between the media and the adventitia contribute to pathological conditions such as aortic aneurysms, thrombosis, or stenosis [24]. Mechanical heterogeneity plays a pivotal role in the pathogenesis of these conditions: Alterations in the isotropic extracellular matrix can lead to vessel dilation, while changes in the anisotropic collagen content can affect overall integrity. Finite element models that account for layer-specific structural and mechanical properties are critical to accurately simulate disease progression, assess rupture risk, and develop targeted interventions [16]. A comprehensive understanding of the interplay between the layers of the aorta can inform strategies for early detection, risk stratification, and tailored therapeutic approaches in the benefit of cardiovascular health.

### Automated model discovery does democratize finite element simulations

For more than half a century, scientists have developed constitutive models for biological tissues [22] and today’s finite element packages offer large libraries of material models to choose from [1, 3, 36, 57, 60]. However, the scientific criteria for appropriate model selection remain highly subjective and prone to user bias. Importantly, the objective of our study is *not* to discover yet another marginally better constitutive model. Instead, our goal is to prototype an *intelligent and automated workflow*—from experiment to simulation— to robustly discover constitutive models from data [33], feed these models directly into a finite element simulation [45], and reliably predict physically meaningful stress and strain profiles. If successful, this new technology could make physics-based simulation more user-friendly, more accessible, and less vulnerable to human error.

## 2 Experiment

We begin by briefly describing our experimental data from the healthy human aorta of a 56-year-old male [40], collected as an intact tube within 24h of death, and stored in saline solution [25]. The sample was cleared, dehydrated in ethanol, and stored in benzyl alcohol-benzyl benzoate, all at room temperature. For the structural characterization, second harmonic generation imaging was used to quantify two microstructural parameters: the collagen fiber angle *α* with respect to the circumferential direction, and the fiber dispersion *κ* [49]. In the circumferential-axial plane, the median collagen fiber angle was *α* = *±*7.00*°* with a fiber dispersion of *κ* = 0.0737 for the media and *α* = *±*66.78*°* with a fiber dispersion of *κ* = 0.0909 for the adventitia [25, 41].

For the mechanical characterization, a squared 20*×*20 mm sample of the media and a cruciform-shaped 35*×*35 mm sample with a squared 5*×*5 mm center testing region of the adventitia were manually separated from the remaining tissue and tested in biaxial extension while submerged in saline solution at 37*°*C. To ensure a homogeneous deformation state, both samples were mounted with the collagen fibers oriented symmetrically with respect to the two loading directions, and loaded at five different stretch ratios, *λ*_cir_ : *λ*_axl_ = {1.20 : 1.10, 1.20 : 1.15, 1.20 : 1.20, 1.15 : 1.20, 1.10 : 1.20}. Tables 1 and 2 summarize the resulting five pairs of datasets, {*λ*_cir_, *σ*_cir_} and {*λ*_axl_, *σ*_axl_}, for the media and for the adventitia [40]. Figure 1 illustrates the circumferential and axial stress-stretch relations of the media, left, and of the adventitia, right, of the 56-year-old healthy human aorta.

**Table 1:** Biaxial testing of human aortic media. Samples are stretched biaxially in the circumferential and axial directions at *λ*_cir_ and *λ*_axl_ at five different stretch ratios. The mean fiber angle is *±*7.00*°* against the circumferential direction. Stresses are reported as *σ*_cir_ and *σ*_axl_, see Figure 1 [25, 40].

| media off-x1.0 |  |  | media off-x1.5 |  |  | media equi-biax |  |  | media off-y1.5 |  |  | media off-y1.0 |  |  |
| --- | --- | --- | --- | --- | --- | --- | --- | --- | --- | --- | --- | --- | --- | --- |
| $\lambda_{\text{cir}} : \lambda_{\text{axl}} = 1.20 : 1.10$ | | | $\lambda_{\text{cir}} : \lambda_{\text{axl}} = 1.20 : 1.15$ | | | $\lambda_{\text{cir}} : \lambda_{\text{axl}} = 1.20 : 1.20$ | | | $\lambda_{\text{cir}} : \lambda_{\text{axl}} = 1.15 : 1.20$ | | | $\lambda_{\text{cir}} : \lambda_{\text{axl}} = 1.10 : 1.20$ | | |
| $\lambda_{\text{cir}}$<br>[-] | $\sigma_{\text{cir}}$<br>[kPa] | $\sigma_{\text{axl}}$<br>[kPa] | $\lambda_{\text{cir}}$<br>[-] | $\sigma_{\text{cir}}$<br>[kPa] | $\sigma_{\text{axl}}$<br>[kPa] | $\lambda_{\text{cir}}$<br>[-] | $\sigma_{\text{cir}}$<br>[kPa] | $\sigma_{\text{axl}}$<br>[kPa] | $\lambda_{\text{cir}}$<br>[-] | $\sigma_{\text{cir}}$<br>[kPa] | $\sigma_{\text{axl}}$<br>[kPa] | $\lambda_{\text{cir}}$<br>[-] | $\sigma_{\text{cir}}$<br>[kPa] | $\sigma_{\text{axl}}$<br>[kPa] |
| 1.000 | 0.00 | 0.00 | 1.000 | 0.00 | 0.00 | 1.000 | 0.00 | 0.00 | 1.000 | 0.00 | 0.00 | 1.000 | 0.00 | 0.00 |
| 1.013 | 1.92 | 0.20 | 1.013 | 1.67 | 0.05 | 1.013 | 1.92 | 0.64 | 1.009 | 1.27 | 0.16 | 1.006 | 0.33 | 0.53 |
| 1.025 | 3.98 | 0.47 | 1.025 | 3.07 | 0.40 | 1.025 | 3.93 | 1.43 | 1.019 | 2.70 | 1.09 | 1.013 | 0.57 | 1.47 |
| 1.038 | 7.14 | 0.89 | 1.038 | 6.01 | 0.79 | 1.038 | 7.57 | 3.85 | 1.028 | 4.82 | 2.82 | 1.019 | 1.76 | 2.56 |
| 1.050 | 11.15 | 2.54 | 1.050 | 10.01 | 2.88 | 1.050 | 12.11 | 6.75 | 1.038 | 7.63 | 5.26 | 1.025 | 3.76 | 4.10 |
| 1.063 | 15.75 | 4.51 | 1.063 | 14.36 | 5.53 | 1.063 | 17.11 | 10.24 | 1.047 | 11.09 | 8.17 | 1.031 | 6.12 | 6.24 |
| 1.075 | 20.86 | 7.05 | 1.075 | 19.07 | 8.76 | 1.075 | 22.56 | 14.43 | 1.056 | 15.21 | 11.19 | 1.038 | 8.72 | 8.77 |
| 1.088 | 26.22 | 9.27 | 1.088 | 24.56 | 12.07 | 1.088 | 28.33 | 18.85 | 1.066 | 19.65 | 14.11 | 1.044 | 11.78 | 11.37 |
| 1.100 | 31.29 | 12.28 | 1.100 | 30.07 | 15.80 | 1.100 | 34.28 | 22.86 | 1.075 | 24.26 | 17.64 | 1.050 | 14.60 | 14.35 |
| 1.113 | 36.77 | 15.20 | 1.113 | 36.02 | 19.45 | 1.113 | 40.44 | 27.00 | 1.084 | 28.87 | 21.54 | 1.056 | 17.62 | 17.54 |
| 1.125 | 42.62 | 18.12 | 1.125 | 42.04 | 23.59 | 1.125 | 47.08 | 32.27 | 1.094 | 33.78 | 25.73 | 1.063 | 20.78 | 20.95 |
| 1.138 | 48.84 | 21.17 | 1.138 | 48.54 | 27.75 | 1.138 | 54.51 | 37.36 | 1.103 | 38.76 | 29.90 | 1.069 | 24.07 | 24.63 |
| 1.150 | 55.32 | 24.44 | 1.150 | 55.49 | 31.61 | 1.150 | 62.11 | 42.29 | 1.113 | 43.77 | 34.20 | 1.075 | 27.44 | 28.40 |
| 1.163 | 62.16 | 27.17 | 1.163 | 62.66 | 35.85 | 1.163 | 70.36 | 47.53 | 1.122 | 49.00 | 38.92 | 1.081 | 30.81 | 32.11 |
| 1.175 | 69.74 | 30.67 | 1.175 | 70.20 | 40.71 | 1.175 | 79.90 | 53.11 | 1.131 | 54.53 | 43.60 | 1.088 | 34.18 | 35.97 |
| 1.188 | 78.37 | 33.93 | 1.188 | 79.51 | 45.23 | 1.188 | 90.40 | 59.34 | 1.141 | 60.95 | 48.59 | 1.094 | 38.11 | 40.00 |
| 1.200 | 89.82 | 39.69 | 1.200 | 90.02 | 51.84 | 1.200 | 101.14 | 66.98 | 1.150 | 69.13 | 55.49 | 1.100 | 43.50 | 45.16 |

**Table 2:** Biaxial testing of human aortic adventitia. Samples are stretched biaxially in the circumferential and axial directions at *λ*_cir_ and *λ*_axl_ at five different stretch ratios. The mean fiber angle is *±*66.78*°* against the circumferential direction. Stresses are reported as *σ*_cir_ and *σ*_axl_, see Figure 1 [25, 40].

| adventitia off-x1.0 |  |  | adventitia off-x1.5 |  |  | adventitia equi-biax |  |  | adventitia off-y1.5 |  |  | adventitia off-y1.0 |  |  |
| --- | --- | --- | --- | --- | --- | --- | --- | --- | --- | --- | --- | --- | --- | --- |
| $\lambda_{\text{cir}} : \lambda_{\text{axl}} = 1.20 : 1.10$ | | | $\lambda_{\text{cir}} : \lambda_{\text{axl}} = 1.20 : 1.15$ | | | $\lambda_{\text{cir}} : \lambda_{\text{axl}} = 1.20 : 1.20$ | | | $\lambda_{\text{cir}} : \lambda_{\text{axl}} = 1.15 : 1.20$ | | | $\lambda_{\text{cir}} : \lambda_{\text{axl}} = 1.10 : 1.20$ | | |
| $\lambda_{\text{cir}}$<br>[-] | $\sigma_{\text{cir}}$<br>[kPa] | $\sigma_{\text{axl}}$<br>[kPa] | $\lambda_{\text{cir}}$<br>[-] | $\sigma_{\text{cir}}$<br>[kPa] | $\sigma_{\text{axl}}$<br>[kPa] | $\lambda_{\text{cir}}$<br>[-] | $\sigma_{\text{cir}}$<br>[kPa] | $\sigma_{\text{axl}}$<br>[kPa] | $\lambda_{\text{cir}}$<br>[-] | $\sigma_{\text{cir}}$<br>[kPa] | $\sigma_{\text{axl}}$<br>[kPa] | $\lambda_{\text{cir}}$<br>[-] | $\sigma_{\text{cir}}$<br>[kPa] | $\sigma_{\text{axl}}$<br>[kPa] |
| 1.000 | 0.00 | 0.00 | 1.000 | 0.00 | 0.00 | 1.000 | 0.00 | 0.00 | 1.000 | 0.00 | 0.00 | 1.000 | 0.00 | 0.00 |
| 1.013 | 0.42 | 0.12 | 1.013 | 0.28 | 0.71 | 1.013 | 1.12 | 0.16 | 1.009 | 0.88 | 0.60 | 1.006 | 0.53 | 0.12 |
| 1.025 | 0.90 | 0.33 | 1.025 | 1.00 | 1.11 | 1.025 | 1.68 | 1.08 | 1.019 | 1.45 | 1.51 | 1.013 | 0.99 | 1.50 |
| 1.038 | 1.51 | 0.94 | 1.038 | 1.59 | 1.63 | 1.038 | 2.52 | 2.28 | 1.028 | 2.22 | 2.57 | 1.019 | 1.53 | 2.00 |
| 1.050 | 2.31 | 1.58 | 1.050 | 2.20 | 2.23 | 1.050 | 3.41 | 3.26 | 1.038 | 3.03 | 3.52 | 1.025 | 2.02 | 2.34 |
| 1.063 | 3.08 | 2.21 | 1.063 | 2.78 | 2.92 | 1.063 | 4.18 | 3.97 | 1.047 | 3.63 | 4.48 | 1.031 | 2.42 | 2.86 |
| 1.075 | 3.73 | 2.80 | 1.075 | 3.62 | 3.73 | 1.075 | 4.95 | 4.73 | 1.056 | 4.07 | 5.45 | 1.038 | 2.90 | 3.84 |
| 1.088 | 4.58 | 3.29 | 1.088 | 4.47 | 4.40 | 1.088 | 5.78 | 5.84 | 1.066 | 4.66 | 6.00 | 1.044 | 3.39 | 4.56 |
| 1.100 | 5.12 | 4.14 | 1.100 | 5.32 | 5.29 | 1.100 | 6.62 | 7.10 | 1.075 | 4.88 | 7.04 | 1.050 | 3.98 | 5.43 |
| 1.113 | 6.28 | 4.66 | 1.113 | 6.28 | 6.24 | 1.113 | 7.48 | 8.64 | 1.084 | 5.52 | 8.90 | 1.056 | 4.40 | 6.11 |
| 1.125 | 6.99 | 5.52 | 1.125 | 7.19 | 7.45 | 1.125 | 8.49 | 10.19 | 1.094 | 5.88 | 10.74 | 1.063 | 5.06 | 7.01 |
| 1.138 | 7.99 | 6.32 | 1.138 | 8.23 | 8.67 | 1.138 | 9.82 | 11.81 | 1.103 | 6.63 | 12.55 | 1.069 | 5.87 | 8.08 |
| 1.150 | 9.28 | 7.01 | 1.150 | 9.48 | 10.05 | 1.150 | 11.26 | 13.47 | 1.113 | 7.82 | 14.37 | 1.075 | 6.70 | 9.47 |
| 1.163 | 10.49 | 8.01 | 1.163 | 10.84 | 11.60 | 1.163 | 12.73 | 15.51 | 1.122 | 9.27 | 16.36 | 1.081 | 7.57 | 11.15 |
| 1.175 | 11.57 | 9.14 | 1.175 | 12.25 | 12.93 | 1.175 | 14.56 | 18.43 | 1.131 | 10.89 | 18.96 | 1.088 | 8.41 | 13.10 |
| 1.188 | 13.10 | 10.05 | 1.188 | 14.22 | 14.78 | 1.188 | 17.23 | 23.00 | 1.141 | 12.74 | 23.06 | 1.094 | 9.82 | 16.50 |
| 1.200 | 15.59 | 12.13 | 1.200 | 17.03 | 17.51 | 1.200 | 20.92 | 29.77 | 1.150 | 17.21 | 32.69 | 1.100 | 11.41 | 23.63 |

**Figure 1:**
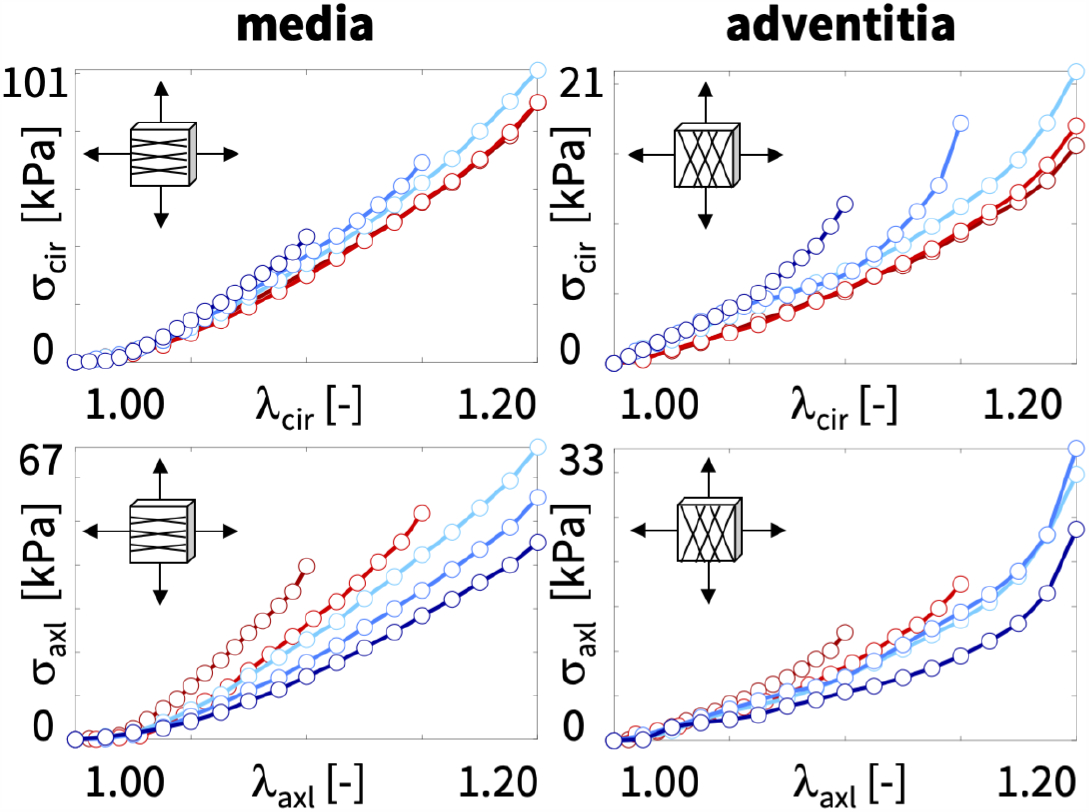
Biaxial testing of human aortic media and adventitia. Samples are stretched biaxially in the circumferential and axial directions at *λ*_cir_ and *λ*_axl_ at five different stretch ratios, from dark red to dark blue. The mean fiber angles of the media and adventitia are *±*7.00*°* and *±*66.78*°* against the circumferential direction. Stresses are reported as *σ*_cir_ and *σ*_axl_, see Tables 1 and 2 [25, 40].

## 3 Model

### Kinematics

During testing, particles ***X*** of the undeformed sample map to particles ***x*** = ***ϕ***(***X***) of the deformed sample via the deformation map ***ϕ***. Its gradient with respect to the undeformed coordinates ***X*** is the deformation gradient, ***F*** = ∇_***X***_***ϕ***. Its spectral representation introduces the principal stretches *λ*_*i*_ and the principal directions ***N***_*i*_ and ***n***_*i*_ in the undeformed and deformed configurations, where ***F*** *·* ***N***_*i*_ = *λ*_*i*_ ***n***_*i*_, and

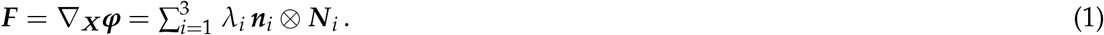

We assume that the vascular tissue has two pronounced fiber directions [21], ***n***_01_ and ***n***_02_, with unit length, ||***n***_01_|| = 1 and ||***n***_02_|| = 1, in the undeformed configuration, which map onto the pronounced directions, ***n***_1_ = ***F n***_01_ and ***n***_2_ = ***F n***_02_, with fiber stretches, ||***n***_1_|| = *λ*_*n*1_ and ||***n***_2_|| = *λ*_*n*2_, in the deformed configuration. We characterize its deformation state through the three principal invariants *I*_1_, *I*_2_, *I*_3_, and six additional invariants *I*_4_, *I*_5_, *I*_6_, *I*_7_, *I*_8_, *I*_9_ [51],

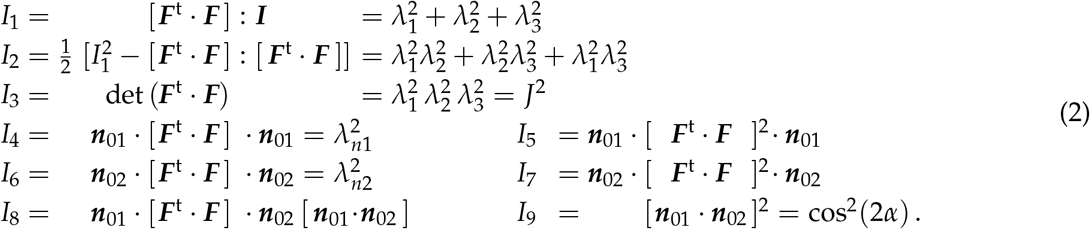

A perfectly incompressible material has a constant Jacobian equal to one, *I*_3_ = *J*^2^ = 1, the ninth invariant is constant by definition, *I*_9_ = const., and the set of independent invariants reduces to seven, *I*_1_, *I*_2_, *I*_4_, *I*_5_, *I*_6_, *I*_7_, *I*_8_.

### Biaxial extension

For the special homogeneous deformation of biaxial extension, we apply stretches *λ*_1_ ≥ 1 and *λ*_2_ ≥ 1 in the *circumferential* and *longitudinal* directions, and adopt the incompressibility condition, 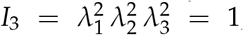 to express the stretch in the *radial* direction, *λ*_3_ = (*λ*_1_ *λ*_2_)^−1^ ≤1. We assume that the fiber pairs, initially oriented at an angle ± *α* to the circumferential direction, ***n***_0_ = [cos(*α*), *±*sin(*α*), 0]^t^, remain symmetric with respect to the stretch directions, such that the deformation remains homogeneous and shear free, and the deformation gradient,

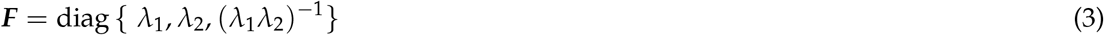

remains diagonal at all times. We now use the the principal stretches *λ*_1_ and *λ*_2_ to express the invariants (2),

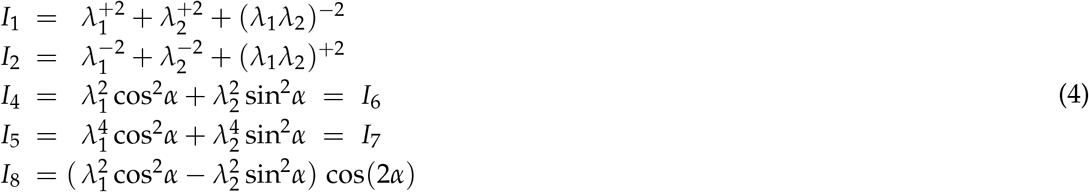

and their derivatives,

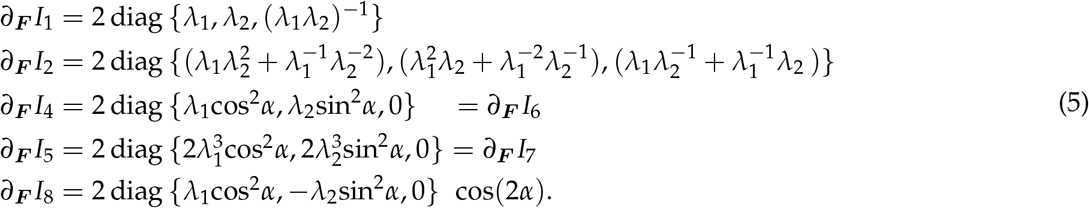

We conclude that the case of biaxial extension probes both fiber directions equally, *I*_4_ = *I*_6_ and *I*_5_ = *I*_7_.

### Constitutive equations

A *hyperelastic* material satisfies the second law of thermodynamics, and its Piola stress ***P*** = ∂*ψ*(***F***)/∂***F*** is the derivative of the free energy *ψ*(***F***) with respect to the deformation gradient ***F***. A *perfectly incompressible* hyperelastic material uses this stress definition modified by a pressure term, −*p* ***F*** ^-t^ [37],

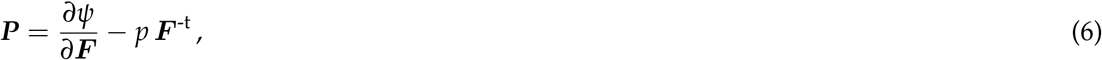

where the hydrostatic pressure, 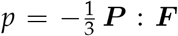, acts as a Lagrange multiplier that we determine from the boundary conditions. We express the free energy function in terms of the seven invariants, *ψ*(*I*_1_, *I*_2_, *I*_4_, *I*_5_, *I*_6_, *I*_7_, *I*_8_), and obtain the following explicit expression for the Piola stress,

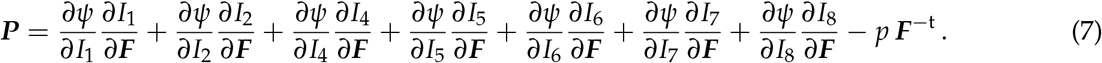

### Biaxial extension

For homogeneous and shear free biaxial extension, the Piola stress ***P*** remains diagonal at all times,

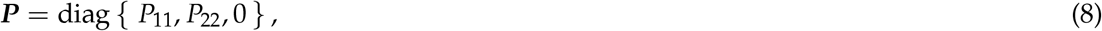

and we can use the zero-normal-stress condition, *P*_33_ = 0, to determine the pressure *p*,

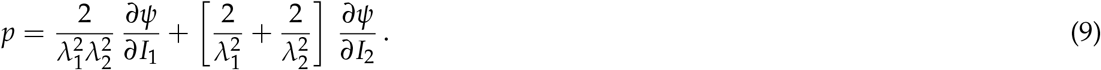

Equation (7) then provides explicit analytical expressions for the Piola stresses *P*_1_ and *P*_2_ in terms of the stretches *λ*_1_ and *λ*_2_, In what follows, we assume that the mechanical behavior of the two fiber families is identical and combine their effects in the fourth and fifth invariants, *I*_4_ and *I*_5_. In addition, we assume that the two fiber families do not interact and drop the eighth invariant *I*_8_ [39]. This results in the following expressions,

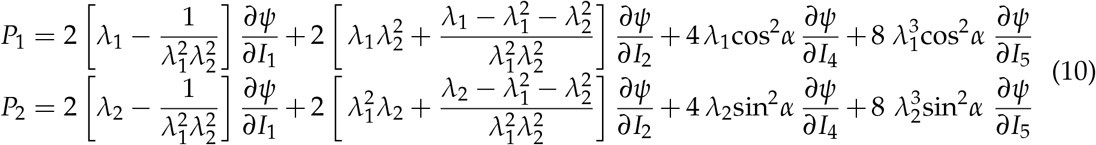

Finally, we translate these nominal stresses *P*_1_ and *P*_2_ into the true stress *σ*_1_ and *σ*_2_,

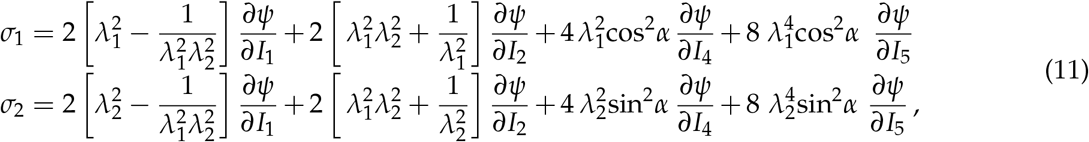

that are reported in the experiment [40].

### Constitutive neural network

To discover the best model and parameters to explain the biaxial testing data, we adopt the concept of constitutive neural networks, a special class of neural networks that satisfy the conditions of thermodynamic consistency, material objectivity, material symmetry, perfect incompressibility, polyconvexity, and physical constraints by design [33].

Figure 2 illustrates our neural network with two hidden layers and eight and sixteen nodes [35]. The first layer generates powers (*∘*) and (*∘*)^2^ of the network input, the four invariants *I*_1_, *I*_2_, *I*_4_, *I*_5_, and the second layer applies the identity, (*∘*) and the exponential function (exp(*∘*)) to these powers. The free energy function of this networks takes the following explicit form,

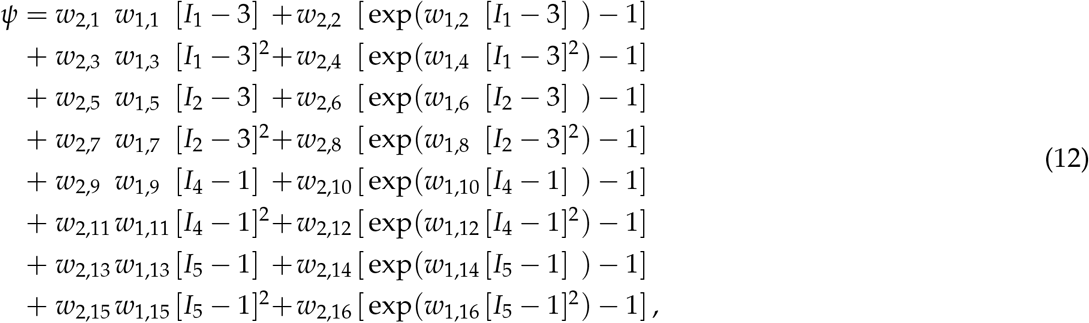

corrected by the pressure term *ψ* = *ψ* − *p* [*J* − 1]. Its derivatives with respect to the four invariants,

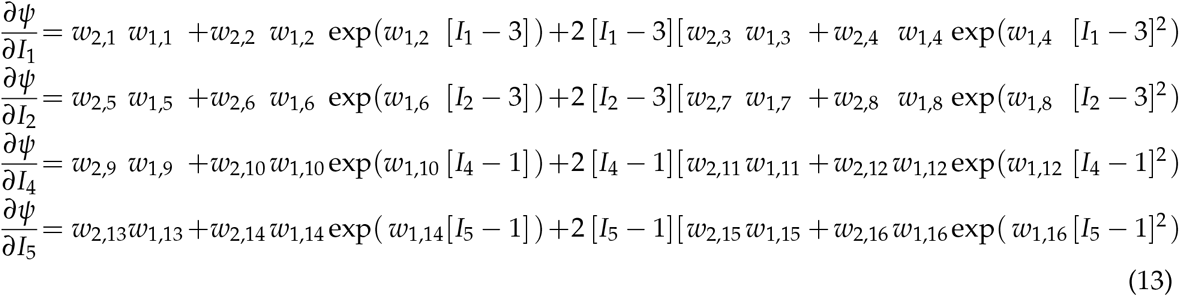

complete the definition of the principal Cauchy stresses in equations (11). The network has two times sixteen weights **w**, which we constraint to always remain non-negative, **w**≥ **0**. We learn the network weights **w** by minimizing a loss function *L* that penalizes the error between model and data. We characterize this error as the mean squared error, the *L*_2_-norm of the difference between the stresses predicted by the network model, *σ*_1_, *σ*_2_, and the experimentally measured stresses, 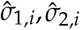 divided by the number of training points *n*_trn_, and add a penalty term, 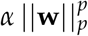 to allow for *L*_*p*_ regularization,

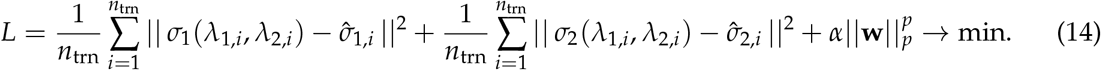

Here *α* ≥ 0 is a non-negative penalty parameter and 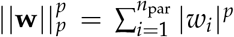 is the *L*_*p*_ norm of the vector of the network weights **w**. We train the network by minimizing the loss function (14) using the ADAM optimizer, a robust adaptive algorithm for gradient-based first-order optimization.

**Figure 2:**
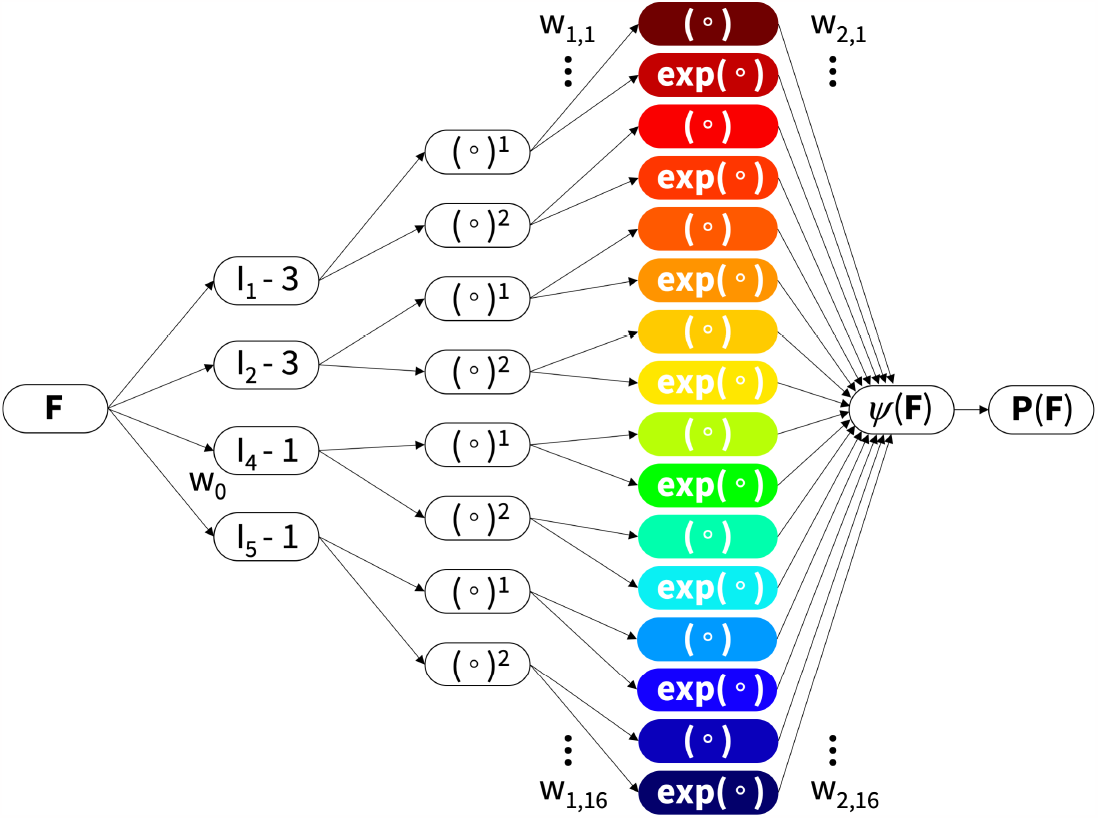
Constitutive neural network. Perfectly incompressible hyperelastic constitutive neural network with two hidden layers to approximate the free-energy function *ψ*(*I*_1_, *I*_2_, *I*_4_, *I*_5_) as a function of the invariants of the deformation gradient ***F*** using sixteen terms. The first layer generates powers (*∘*) and (*∘*)^2^ of the network input and the second layer applies the identity (*∘*) and exponential function (exp(*∘*)) to these powers.

### Universal material subroutine

To seamlessly integrate our discovered model and parameters into a simulation, we create a universal material subroutine [45]. This subroutine operates on the integration point level of the finite element analysis and translates the local deformation, for example in the form of the deformation gradient ***F***, into the current stress, for example the Piola stress ***P*** [1]. We reformulate the free energy function *ψ* from equation (12) as the sum of all *k* nodes of the final hidden layer,

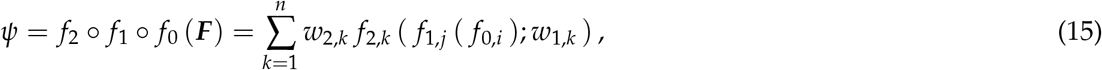

where *f*_2_, *f*_1_, *f*_0_ are the nested activation functions associated with the second, first, and zeroth layers,

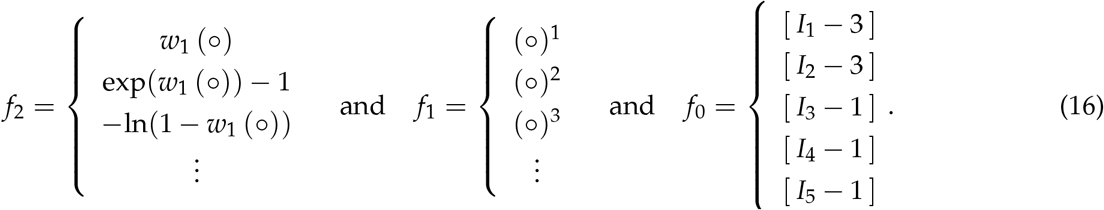

Here *f*_0_ maps the deformation gradient ***F*** onto a set of invariants, [*I*_1_ − 3], [*I*_2_ − 3], [*I*_3_ − 1], [*I*_4_ − 1], [*I*_5_ − 1], *f*_1_ raises these invariants to the first, second, or any higher order powers, (∘)^1^, (∘)^2^, (∘)^3^, and *f*_2_ applies the identity, exponential, or natural logarithm, (∘), (exp(∘) 1), (ln(1 (∘))), or any other thermodynamically admissible function to these powers. The material subroutine calculates the Piola stress following equation (13),

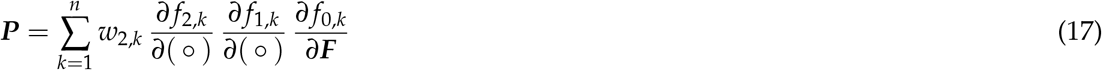

in terms of the first derivatives of the activation functions *f*_2_ and *f*_1_,

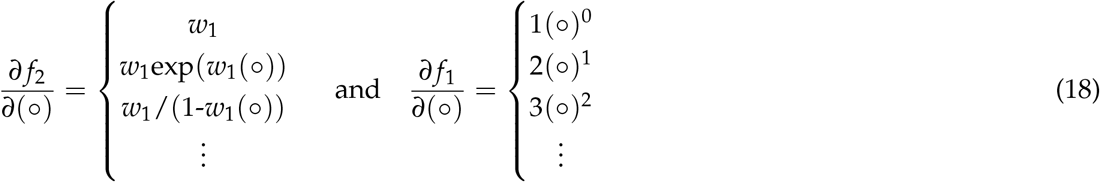

and the tensor basis, ∂ *f*_0,*k*_/∂***F*** = ∂*I*_*k*_/∂***F***. In implicit finite element algorithms with a global Newton Raphson iteration, the material subroutine also calculates the tangent moduli,

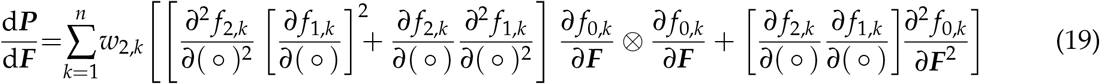

in terms of the second derivatives of the activation functions *f*_2_ and *f*_1_,

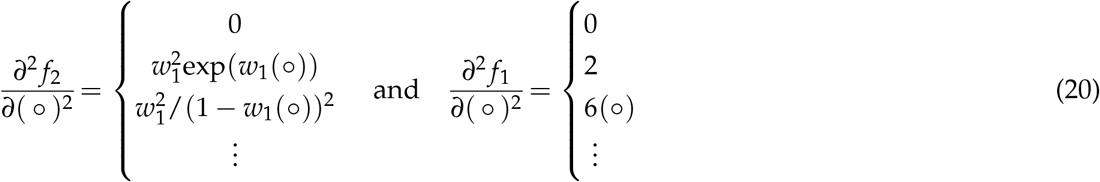

and the tensor basis, ∂^2^ *f*_0,*k*_/∂***F***^2^ = ∂^2^ *I*_*k*_/∂***F***⊗∂***F***. We translate our discovered model into a modular universal material subroutine within the Abaqus finite element analysis software suite [1]. We leverage the `UANISOHYPER_INV` subroutine to introduce our strain energy function (12) or (16) in terms of the discovered pairs of network weights and activation functions. Our universal material subroutine uses the strain energy density, `UA(1)` = *ψ*, and its first and second derivatives, `UI1(NINV)` = ∂*ψ*/∂*I*_*i*_, and `UI2(NINV`^∗^`(NINV+1)/2)`= ∂^2^*ψ*/∂*I*_*i*_∂*I*_*j*_, with respect to the invariants. Following the Abaqus convention, we introduce an array of generalized invariants, 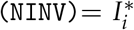 with *i* = 1, …,`NINV`, where `NINV` is the total number of isotropic and anisotropic invariants. In our case, for a material with two fiber families, ***n***_0_ = [cos(*α*), ± sin(*α*), 0]^t^, we introduce four additional invariants, *I*_4_, *I*_5_, *I*_6_, *I*_7_, where *I*_4_, *I*_6_ and *I*_5_, *I*_7_ share the same parameters [1].

Algorithm 1 illustrates the `UANISOHYPER_INV` pseudocode to compute the arrays, `UA(1)`, `UI1(NINV)`, `UI2(NINV` ^∗^`(NINV+1)/2)`, at the integration point level during a finite element analysis. First, we initialize all relevant arrays and read the activation functions *k f*_1,*k*_ and *k f*_2,*k*_ and weights *w*_1,*k*_ and *w*_2,*k*_ of the *n* color-coded nodes of our network in Figure 2 from our user-defined parameter table `UNIVERSAL_TAB`. Then, for each node, we evaluate its row in the parameter table `UNIVERSAL_TAB` and additively update the strain energy density function and its first and second derivatives, `UA`, `UI1`, `UI2`. Algorithm 2 summarizes the additive update of the free energy and its first and second derivatives, `UA`, `UI1`, `UI2`, within the universal material subroutine `uCANN`. Algorithms 3 and 4 summarize the two subroutines `uCANN_h1` and `uCANN_h2` that evaluate the first and second network layers for each network node with its activation functions and weights.

#### Algorithm 1

Pseudocode for universal material subroutine `UANISOHYPER_INV`

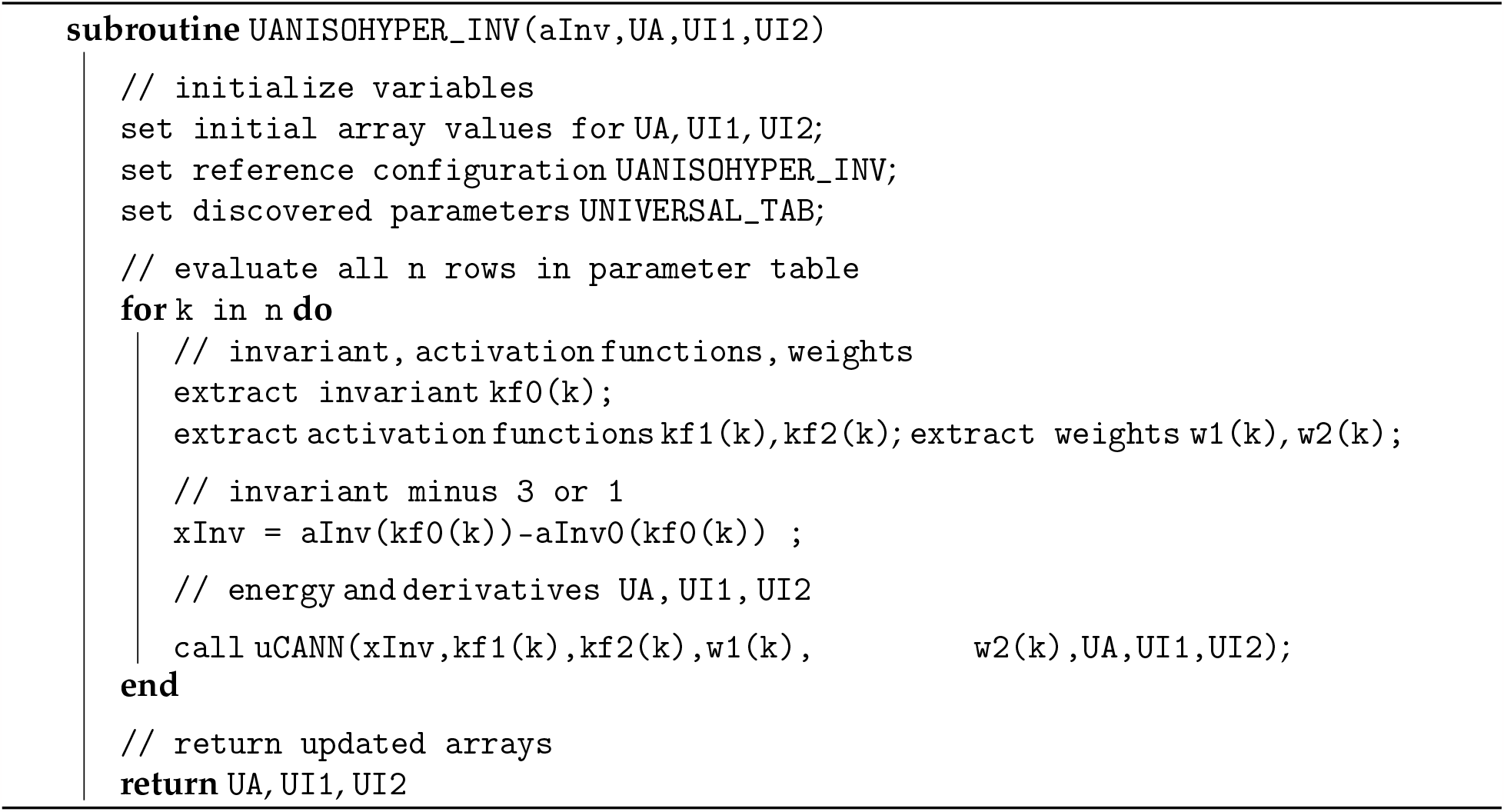

#### Algorithm 2

Pseudocode to update energy and its derivatives `UA, UI1, UI2`

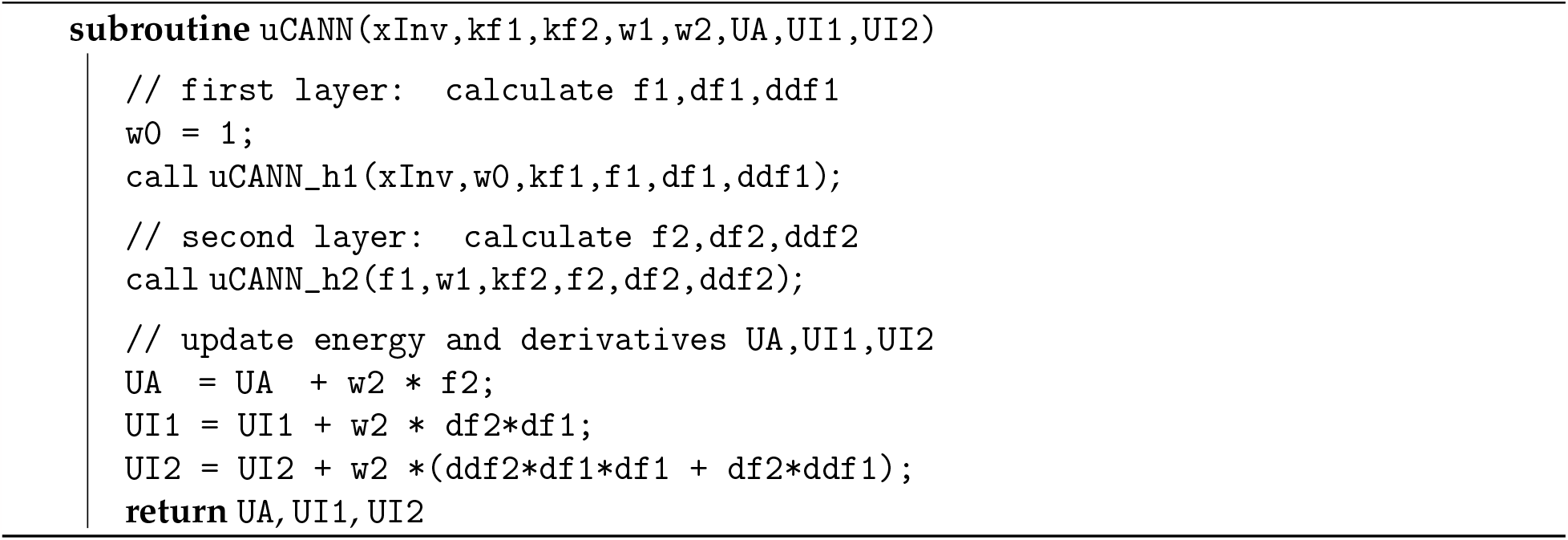

#### Algorithm 3

Pseudocode to evaluate output of first network layer `f,df,ddf`

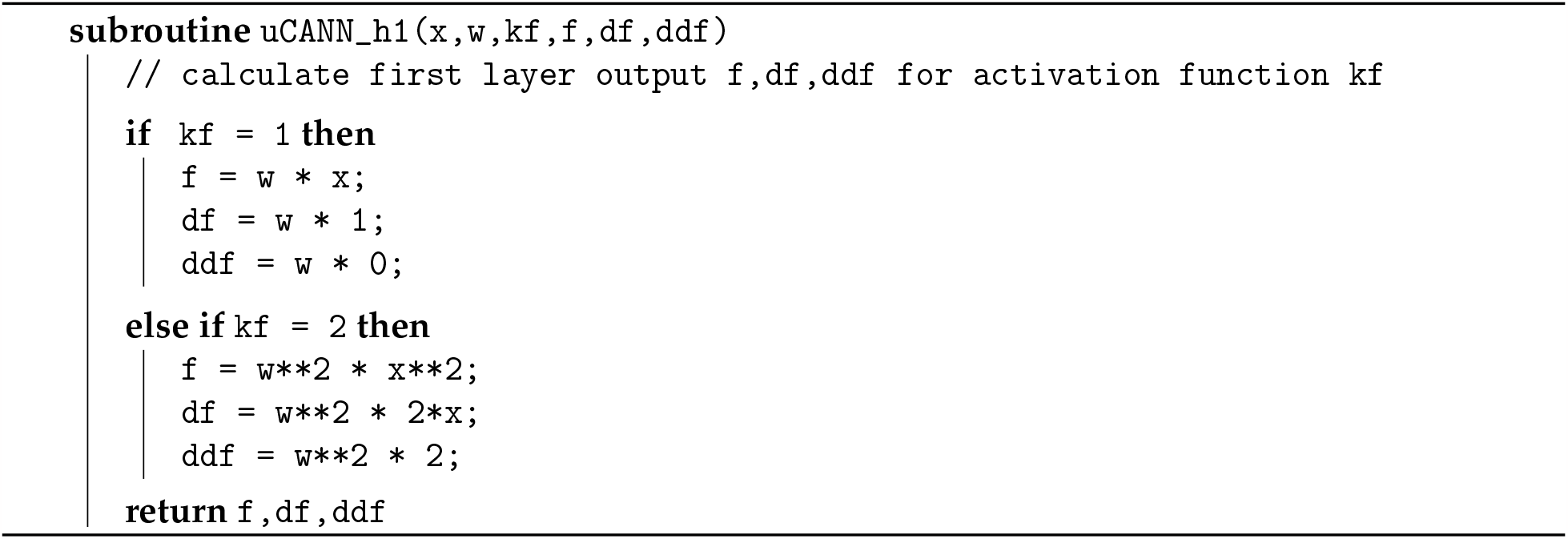

#### Algorithm 4

Pseudocode to evaluate output of second network layer `f,df,ddf`

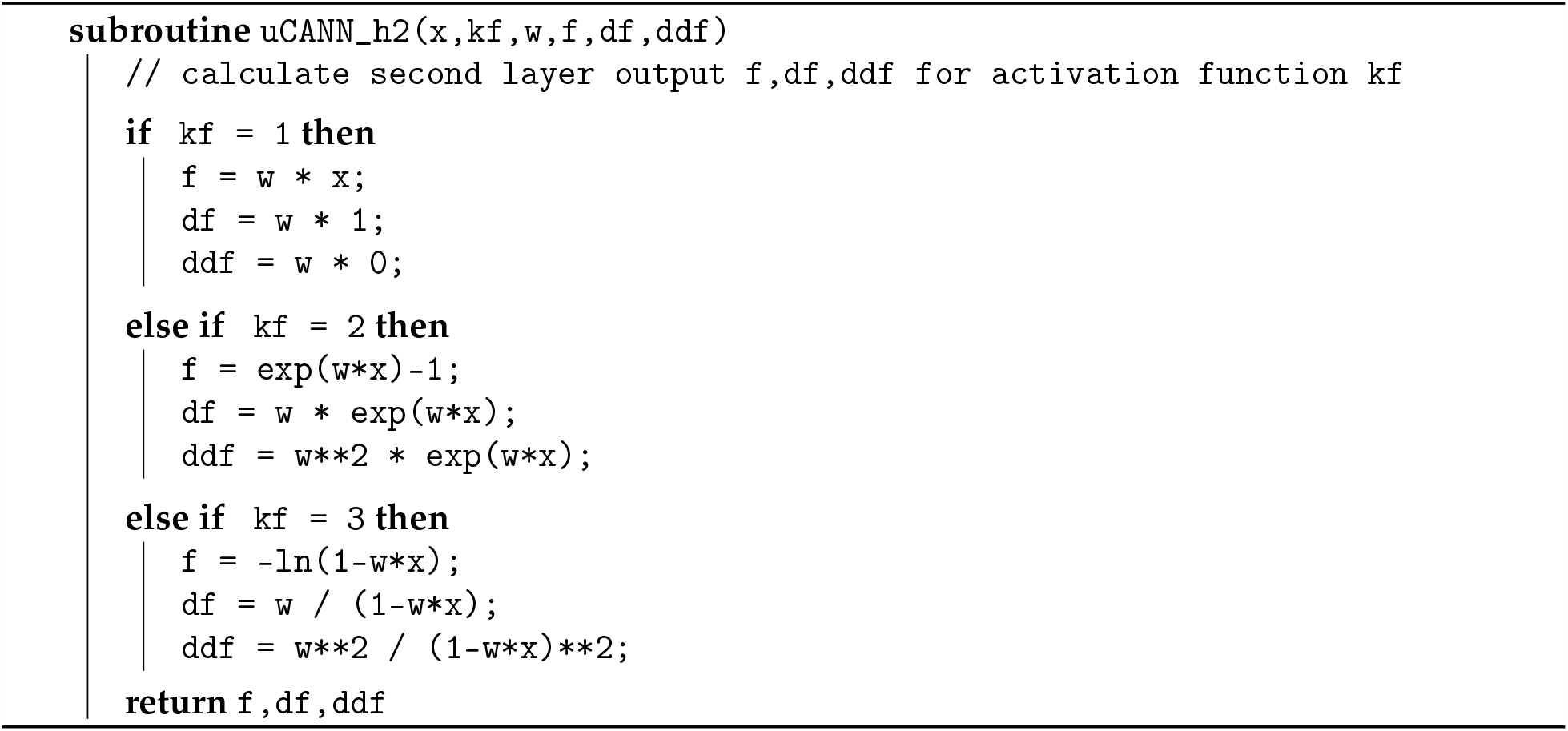

### Finite element simulation

We implement the universal material subroutine in Abaqus FEA, and make it publicly available on Github. To integrate it into a finite element simulation, we need to define our discovered model and parameters in a parameter table [1]. Each row of this table represents one of the color-coded nodes in Figure 1 and consists of five terms: an integer `kf0` that defines the index of the pseudo-invariant `xInv`, two integers `kf1` and `kf2` that define the indices of the first- and second-layer activation functions, and two float values `w1` and `w2` that define the weights of the first and second layers. We declare this input format using the parameter table type definition in the `UNIVERSAL_PARAM_TYPES.INC` file.

~~~
   *PARAMETER TABLE TYPE, name=“UNIVERSAL_TAB”, parameters = 5
   INTEGER, ,”index pseudo-invariant, kf0,o”
   INTEGER, ,”index 1st activ function, kf1,o”
   INTEGER, ,”index 2nd activ function, kf2,o”
   FLOAT, ,”weight 1st hidden layer, w1,o”
   FLOAT, ,”weight 2nd hidden layer, w2,o”
~~~

Within Abaqus FEA, we include the parameter table type definition using

~~~
   *INCLUDE, INPUT=UNIVERSAL_PARAM_TYPES.INC
~~~

at the beginning of the input file. We activate our user-defined material model through the command

~~~
   *ANISOTROPIC HYPERELASTIC, USER, FORMULATION=INVARIANT
~~~

followed by the discovered parameters. From the constitutive neural network in Figure 2, we obtain sixteen entries for the parameter table, four for each isotropic invariant, *I*_1_ and *I*_2_, and four for each anisotropic invariant, *I*_4_ and *I*_5_, associated with the first fiber family, ***n***_0_ = [cos(*α*), + sin(*α*), 0]^t^. We add eight entries, four for each anisotropic invariant, *I*_6_ and *I*_7_, indexed in Abaqus as invariants `8` and `9`, associated with the second fiber family, ***n***_0_ = [cos(*α*), −sin(*α*), 0]^t^, with the same parameters as *I*_4_ and *I*_5_. The header and the twenty-four lines of our parameter table take the following format,

~~~
   *PARAMETER TABLE, TYPE=“UNIVERSAL_TAB”
   1,1,1,w_1,1_,w_2,1_       1,1,2,w_1,2_,w_2,2_      1,2,1,w_1,3_,w_2,3_     1,2,2,w_1,4_,w_2,4_
   2,1,1,w_1,5_,w_2,5_       2,1,2,w_1,6_,w_2,6_      2,2,1,w_1,7_,w_2,7_     2,2,2,w_1,8_,w_2,8_
   4,1,1,w_1,9_,w_2,9_       4,1,2,w_1,10_,w_2,10_     4,2,1,w_1,11_,w_2,11_   4,2,2,w_1,12_,w_2,12_
   5,1,1,w_1,13_,w_2,13_      5,1,2,w_1,14_,w_2,14_     5,2,1,w_1,15_,w_2,15_   5,2,2,w_1,16_,w_2,16_
   8,1,1,w_1,9_,w_2,9_       8,1,2,w_1,10_,w_2,10_     8,2,1,w_1,11_,w_2,11_   8,2,2,w_1,12_,w_2,12_
   9,1,1,w_1,13_,w_2,13_      9,1,2,w_1,14_,w_2,14_     9,2,1,w_1,15_,w_2,15_   9,2,2,w_1,16_,w_2,16_
~~~

The first index of each row selects between the first, second, fourth, fifth, sixth, and seventh invariants, *I*_1_, *I*_2_, *I*_4_, *I*_5_, *I*_6_, *I*_7_, the second index raises them to linear or quadratic powers, (∘)^1^, (∘)^2^, and the third index selects between the identity or the exponential function, (∘), (exp(∘)−1). For brevity, we can simply exclude terms with zero weights from the list.

## 4 Results

To demonstrate how we can translate information seamlessly from experiment to simulation, we perform three types of examples: First, we discover the best model and parameters to explain the experimental data with a limited number of model terms. We discover the *best-in-class one- and two-term models*, interpret their terms, and discuss their model parameters. For the four best-in-class two-term models, we illustrate the fit to the data, and perform a direct comparison with the widely used classical Holzapfel model. Second, we discover the *best model and parameters* to explain the data, but now without restricting the number of terms. We demonstrate how to embed the model into our universal material subroutine, and validate its implementation by comparing its finite element simulations against the experimental data and against the stress plots from our initial model discovery. Third, we predict the diastolic and systolic wall stretches and stresses across a human aortic arch, and compare the simulations with our newly discovered model against the classical Holzapfel model. We illustrate how to parameterize the two models and discuss their similarities and differences, locally at the integration point level and globally at the structural level.

### Discovering the best-in-class models

First, to gain a better intuition of our data, we discover the best families of models with a limited number of terms. In the most general sense, our sixteen-node network in Figure 2 introduces the sixteen-term model in equation (12) parameterized in terms of sixteen pairs of weights, *{w*_1,*∘*_, *w*_2,*∘*_*}*. In the most naive approach, we could test all possible models. From combinatorics, we know that this is a total of 2^16^ − 1 = 65, 535 models, 16 with a single term, 120 with two, 560 with three, 1,820 with four, 4,368 with five, 8,008 with six, 11,440 with seven, 12,870 with eight, 11,440 with nine, 8,008 with ten, 4,368 with eleven, 1,820 with twelve, 560 with thirteen, 120 with fourteen, 16 with fifteen, and 1 with all sixteen terms. To understand the relevance of these sixteen terms, we begin with a simplified analysis that constrains the number of non-zero terms to either one or two [38]. We train the network in Figure 2 by minimizing the loss function (14) with the stress definitions (11) using the biaxial test data of the media and adventitia in Tables 1 and 2, and explicitly set the weights of the remaining terms to zero.

Figure 3 summarizes the discovery of the best-in-class one- and two-term models for the human aortic media and adventitia in two 16*×*16 heat maps. Terms 1 through 8 are associated with the isotropic invariants *I*_1_ and *I*_2_, terms 9 through 16 are associated with the anisotropic invariants *I*_4_ and *I*_5_. The squares on the diagonale indicate the goodness of fit of the 16 one-term models for the media and the adventitia. All other squares indicate the goodness of fit of the 120 two-term models. The color code represents the remaining loss after training, and is a measure for the goodness of fit of each model. The best-in-class models are the models with the lowest remaining loss, highlighted in dark blue. At first glance, we observe four distinct blocks, the iso-iso block in the upper left, the aniso-aniso block in the lower right, and the iso-aniso blocks in the upper right and lower left. The color code confirms our intuition, that a combination of two isotropic or two anisotropic terms does not provide a good explanation of the data. Instead, the best-in-class models with the lowest remaining loss and the dark blue colors are all located in the iso-aniso blocks.

**Figure 3:**
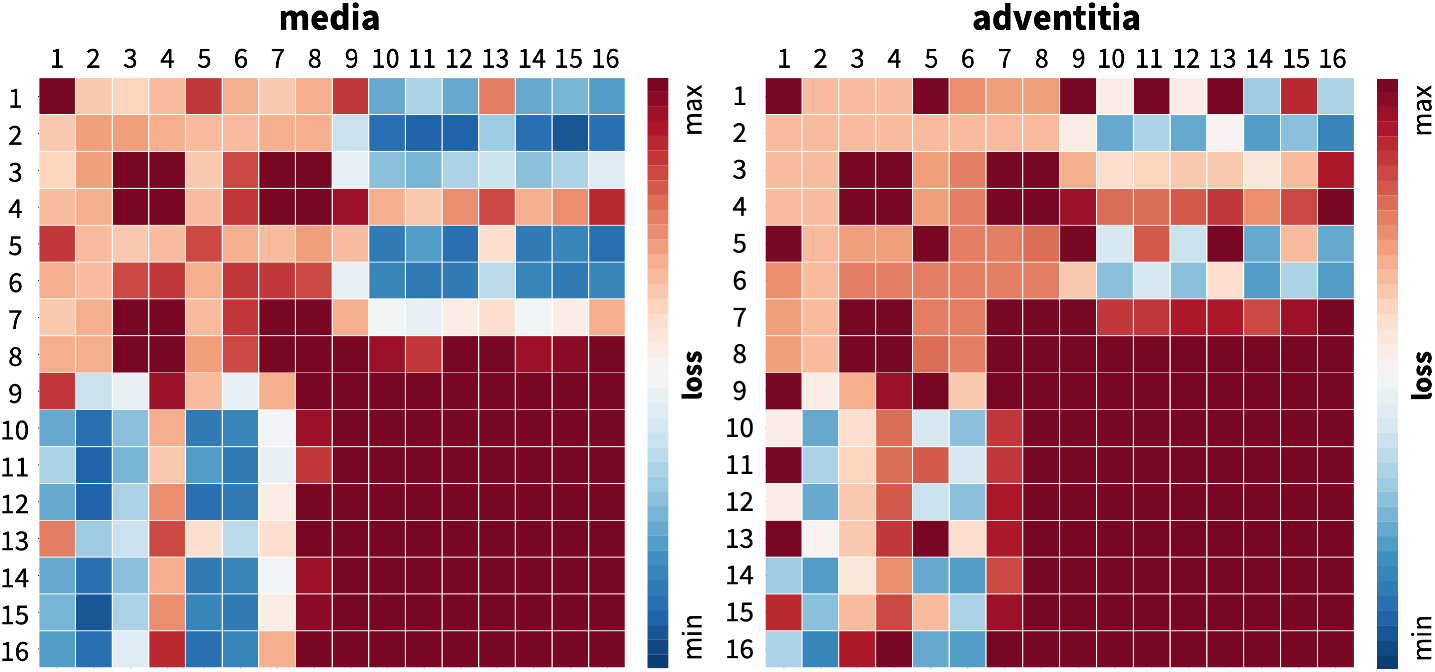
Discovering the best-in-class models. Best-in-class one- and two-term models of the media and adventitia. Remaining loss of the 16 one-term models and 120 two-term models of the constitutive neural network from Figure 2, trained with all ten datasets from Table 1 for the media, left, and Table 2 for the adventitia, right. Terms 1 through 8 are associated with the isotropic invariants *I*_1_ and *I*_2_, terms 9 through 16 are associated with the anisotropic invariants *I*_4_ and *I*_5_. Squares on the diagonale indicate the losses of the 16 one-term models, all other squares indicate the losses of the 120 two-term models. Best-in-class models are the models with the lowest remaining loss, highlighted in dark blue.

### Best-in-class one-term models

The squares on the diagonales of Figure 3 indicate the goodness of fit of the 16 one-term models for the media and the adventitia. Table 3 summarizes the four best-in-class one-term models: the exponential linear first invariant Demiray model [9], the linear second invariant Blatz Ko model [4], the exponential linear second invariant model, and the linear first invariant neo Hooke model [59]. Each block summarizes the constitutive model, the input to the universal material subroutine, their parameterizations for the media, top, and adventita, bottom, and their overall ranking, right. Since our constitutive neural network uses parameters with a clear physical interpretation, we can translate the network weights into the classical shear modulus *μ*, the stiffness-like parameter *a*, and the unitless exponential weighting factor *b*. From comparing the discovered parameters for both tissue types across all four models, we conclude that the media, in each top row, is about three to four times stiffer than the adventitia, in each bottom row. Interestingly, the exponential first invariant Demiray model [9] is the best of all sixteen models, both for the media and adventitia. The linear second invariant Blatz Ko model [4] is the second best model for the media, and the third best for the adventitia. Strikingly, the widely used linear first invariant neo Hooke model [59] is not among the three best-in-class one-term models, neither for the media nor for the adventitia.

**Table 3:** Best-in-class one-term models. Models and parameters of the constitutive neural network from Figure 2, trained with data from Table 1 for the media and Table 2 for the adventita. The four models are the best-in-class one-term models from Figure 2. Each block summarizes the constitutive model, the input to the universal material subroutine, their parameterizations for the media, top, and adventita, bottom, and their overall ranking.

| node 2 - exponential linear first invariant - Demiray |  |  |
| --- | --- | --- |
| $\psi = \frac{1}{2}a/b[\exp(b[I_1 - 3]) - 1]$ | $1, 1, 2, w_{1,2}, w_{2,2}$ | |
| $a = 30.46 \text{ kPa}, b = 3.09$ | $w_{1,2} = 3.09, w_{2,2} = 4.93 \text{ kPa}$ | #1 |
| $a = 9.65 \text{ kPa}, b = 2.59$ | $w_{1,2} = 2.59, w_{2,2} = 1.87 \text{ kPa}$ | #1 |
| node 5 - linear second invariant - Blatz Ko |  |  |
| $\psi = \frac{1}{2}\mu [I_2 - 3]$ | $2, 1, 1, w_{1,5}, w_{2,5}$ | |
| $\mu = 45.93 \text{ kPa}$ | $w_{1,5} = 22.96, w_{2,5} = 1.00 \text{ kPa}$ | #2 |
| $\mu = 12.67 \text{ kPa}$ | $w_{1,5} = 6.34, w_{2,5} = 1.00 \text{ kPa}$ | #3 |
| node 6 - exponential linear second invariant |  |  |
| $\psi = \frac{1}{2}a/b[\exp(b[I_2 - 3]) - 1]$ | $2, 1, 2, w_{1,6}, w_{2,6}$ | |
| $a = 24.55 \text{ kPa}, b = 2.25$ | $w_{1,6} = 2.25, w_{2,6} = 5.46 \text{ kPa}$ | #3 |
| $a = 8.52 \text{ kPa}, b = 1.57$ | $w_{1,6} = 1.57, w_{2,6} = 2.71 \text{ kPa}$ | #2 |
| node 1 - linear first invariant - neo Hooke |  |  |
| $\psi = \frac{1}{2}\mu [I_1 - 3]$ | $1, 1, 1, w_{1,1}, w_{2,1}$ | |
| $\mu = 58.24 \text{ kPa}$ | $w_{1,1} = 29.12, w_{2,1} = 1.00 \text{ kPa}$ | #4 |
| $\mu = 16.04 \text{ kPa}$ | $w_{1,1} = 8.02, w_{2,1} = 1.00 \text{ kPa}$ | #5 |

### Best-in-class two-term models of the media

All squares that are not located on the diagonale of Figure 3 illustrate the goodness of fit of the 120 two-term models. Notably, for the media, the four best-in-class two-term models are all located in the second row and column of Figure 3, left. Table 4 summarizes the four best-in-class two-term models for the media. They all contain the isotropic exponential linear first invariant Demiray term [9] from the best-in-class one-term model, combined with an anisotropic term: the quadratic fifth invariant term, the quadratic fourth invariant term, the exponential quadratic fourth invariant term, or the exponential quadratic fifth invariant term. For comparison, Table 4 also reports the classical two-term Holzapfel model [21] that contains the isotropic linear first invariant term and the anisotropic exponential quadratic fourth invariant term. Each block of the table summarizes the constitutive model, the input to the universal material subroutine, and their parameterizations.

**Table 4:**
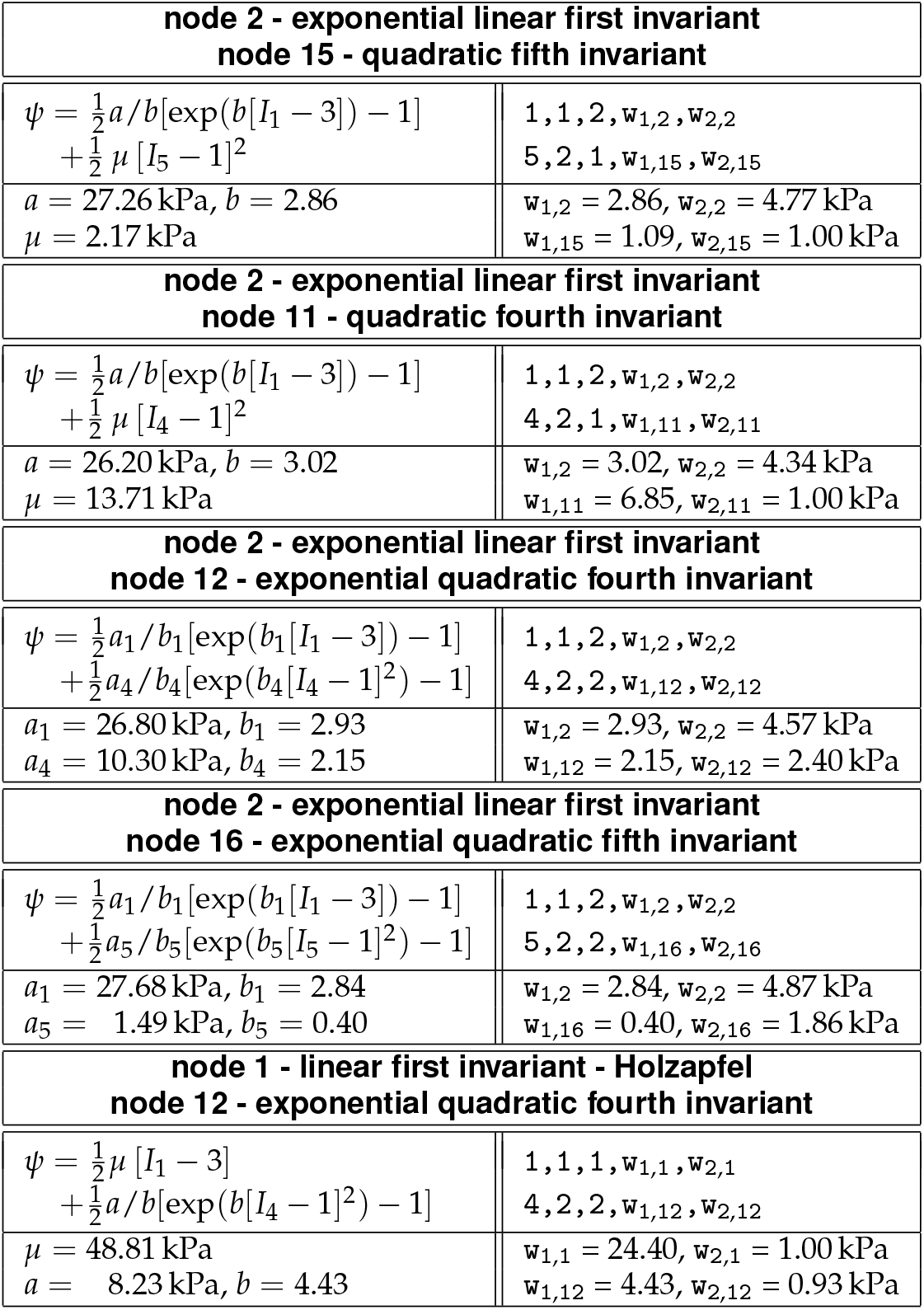
Best-in-class two-term models of the media. Models and parameters of the constitutive neural network from Figure 2, trained with all ten datasets from Table 1 simultaneously. The first four models are the best-in-class two-term models from Figure 2, left; the fifth model is the classical Holzapfel model. Each block summarizes the constitutive model, the input to the material subroutine, and their parameters.

| **node 2 - exponential linear first invariant**<br>**node 15 - quadratic fifth invariant** |  |
| --- | --- |
| $\psi = \frac{1}{2}a/b[\exp(b[I_1 - 3]) - 1] + \frac{1}{2}\mu[I_5 - 1]^2$ | $1, 1, 2, w_{1,2}, w_{2,2}$<br>$5, 2, 1, w_{1,15}, w_{2,15}$ |
| $a = 27.26 \text{ kPa}, b = 2.86$<br>$\mu = 2.17 \text{ kPa}$ | $w_{1,2} = 2.86, w_{2,2} = 4.77 \text{ kPa}$<br>$w_{1,15} = 1.09, w_{2,15} = 1.00 \text{ kPa}$ |
| **node 2 - exponential linear first invariant**<br>**node 11 - quadratic fourth invariant** |  |
| $\psi = \frac{1}{2}a/b[\exp(b[I_1 - 3]) - 1] + \frac{1}{2}\mu[I_4 - 1]^2$ | $1, 1, 2, w_{1,2}, w_{2,2}$<br>$4, 2, 1, w_{1,11}, w_{2,11}$ |
| $a = 26.20 \text{ kPa}, b = 3.02$<br>$\mu = 13.71 \text{ kPa}$ | $w_{1,2} = 3.02, w_{2,2} = 4.34 \text{ kPa}$<br>$w_{1,11} = 6.85, w_{2,11} = 1.00 \text{ kPa}$ |
| **node 2 - exponential linear first invariant**<br>**node 12 - exponential quadratic fourth invariant** |  |
| $\psi = \frac{1}{2}a_1/b_1[\exp(b_1[I_1 - 3]) - 1] + \frac{1}{2}a_4/b_4[\exp(b_4[I_4 - 1]^2) - 1]$ | $1, 1, 2, w_{1,2}, w_{2,2}$<br>$4, 2, 2, w_{1,12}, w_{2,12}$ |
| $a_1 = 26.80 \text{ kPa}, b_1 = 2.93$<br>$a_4 = 10.30 \text{ kPa}, b_4 = 2.15$ | $w_{1,2} = 2.93, w_{2,2} = 4.57 \text{ kPa}$<br>$w_{1,12} = 2.15, w_{2,12} = 2.40 \text{ kPa}$ |
| **node 2 - exponential linear first invariant**<br>**node 16 - exponential quadratic fifth invariant** |  |
| $\psi = \frac{1}{2}a_1/b_1[\exp(b_1[I_1 - 3]) - 1] + \frac{1}{2}a_5/b_5[\exp(b_5[I_5 - 1]^2) - 1]$ | $1, 1, 2, w_{1,2}, w_{2,2}$<br>$5, 2, 2, w_{1,16}, w_{2,16}$ |
| $a_1 = 27.68 \text{ kPa}, b_1 = 2.84$<br>$a_5 = 1.49 \text{ kPa}, b_5 = 0.40$ | $w_{1,2} = 2.84, w_{2,2} = 4.87 \text{ kPa}$<br>$w_{1,16} = 0.40, w_{2,16} = 1.86 \text{ kPa}$ |
| **node 1 - linear first invariant - Holzapfel**<br>**node 12 - exponential quadratic fourth invariant** |  |
| $\psi = \frac{1}{2}\mu[I_1 - 3] + \frac{1}{2}a/b[\exp(b[I_4 - 1]^2) - 1]$ | $1, 1, 1, w_{1,1}, w_{2,1}$<br>$4, 2, 2, w_{1,12}, w_{2,12}$ |
| $\mu = 48.81 \text{ kPa}$<br>$a = 8.23 \text{ kPa}, b = 4.43$ | $w_{1,1} = 24.40, w_{2,1} = 1.00 \text{ kPa}$<br>$w_{1,12} = 4.43, w_{2,12} = 0.93 \text{ kPa}$ |

Figure 4 illustrates the performance of the four best-in-class two-term models for the media from Figure 2, left, summarized in Table 4, and for comparison, the classical Holzapfel model [21]. The circles represent the equibiaxial testing data from Table 1. The reported loss quantifies the goodness of fit for a simultaneous training with all all ten stress-stretch pairs. The color coded regions highlight the contributions of the individual model terms to the circumferential and axial stresses, *σ*_cir_ and *σ*_axl_, as functions of stretches *λ*_cir_ and *λ*_axl_. The red regions represent the isotropic exponential linear first invariant term. The blue, green, turquoise, and dark blue regions represent the anisotropic fourth and fifth invariant terms. With a median collagen fiber orientation of 7.00*°*, the fibers in the media are almost aligned with the circumferential direction. This implies that the axial direction, bottom, only sees the red isotropic response, while the circumferential direction, top, sees a superposition of both, the red isotropic and the green-to-blue anisotropic responses. For the sake of compactness, we only display the equibiaxial response, but note that the other four curves provide an equally good fit to the experimental data. The classical Holzapfel model [21] in Figure 4, right, combines the isotropic linear first invariant term in dark red and the anisotropic exponential quadratic fourth invariant term in turquoise. While it performs well in the circumferential direction, top right, its linear isotropic term is incapable of capturing the nonlinear isotropic matrix behavior in the axial direction, bottom right. Its loss is about three times higher than the loss of the discovered best-in-class two-term-model, Figure 4, left.

**Figure 4:**
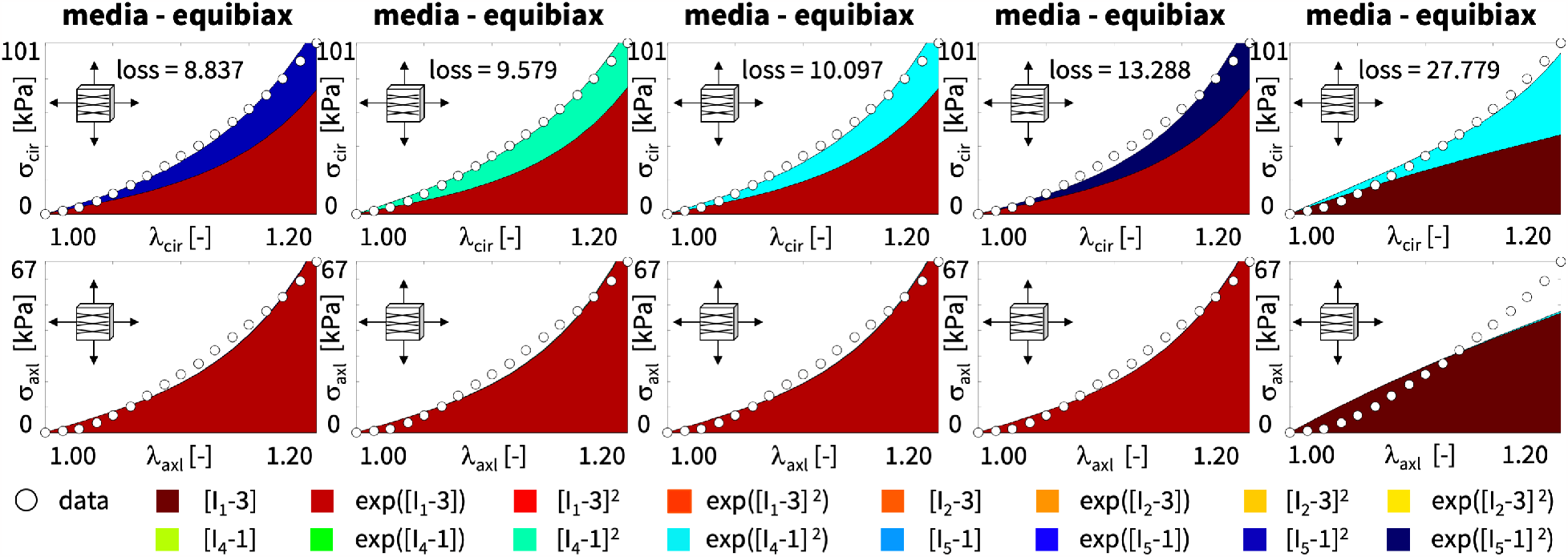
Best-in-class two-term models of the media. True stresses *σ*_cir_ and *σ*_axl_ as functions of stretches *λ*_cir_ and *λ*_axl_ for the constitutive neural network from Figure 2, trained with all ten datasets from Table 1 simultaneously. The first four columns illustrate the best-in-class two-term models from Figure 2, left; the right column illustrates the Holzapfel model [21] for comparison. Circles represent the equibiaxial testing data from Table 1. Color-coded regions represent the discovered model terms. The remaining loss indicates the quality of the overall fit.

### Best-in-class two-term models of the adventitia

All squares that are not located on the diagonale of Figure 3 illustrate the goodness of fit of the 120 two-term models. Interestingly, for the adventitia, the four best-in-class two-term models are located in the second and sixth rows and columns of Figure 3, right. Table 5 summarizes the four best-in-class two-term models for the adventitia. They contain the isotropic exponential linear first or second invariant term from the best-in-class one-term models, combined with the anisotropic exponential linear or quadratic fifth invariant term. For comparison, Table 5 also reports the classical two-term Holzapfel model [21]. Figure 5 illustrates the performance of the four best-in-class two-term models for the adventitia from Figure 2, right, summarized in Table 5, and for comparison, the classical Holzapfel model [21]. The circles represent the equibiaxial testing data from Table 2. The red and light orange regions represent the isotropic exponential linear first and second invariant terms. The blue and dark blue regions represent the anisotropic exponential linear and quadratic fifth invariant terms. With a median collagen fiber orientation of 66.78*°*, the fibers in the adventitia are much closer to the axial direction than for the media. As a result, both circumferential and axial directions see the red and and orange isotropic response and the blue anisotropic response, with a pronounced anisotropy in the axial stresses, bottom. Similar to the media, Figure 4, the classical Holzapfel model [21] for the adventitia, Figure 5, right, has a loss that is about three times higher than the loss of the discovered best-in-class two-term-model, Figure 5, left.

**Table 5:** Best-in-class two-term models of the adventitia. Models and parameters of the constitutive neural network from Figure 2, trained with all ten datasets from Table 2 simultaneously. The first four models are the best-in-class two-term models from Figure 2, right; the fifth model is the classical Holzapfel model. Each block summarizes the constitutive model, the input to the material subroutine, and their parameters.

|  |  |
| --- | --- |
| **node 2 - exponential linear first invariant**<br>**node 16 - exponential quadratic fifth invariant** |  |
| $\psi = \frac{1}{2}a_1/b_1[\exp(b_1[I_1 - 3]) - 1]$<br>$+ \frac{1}{2}a_5/b_5[\exp(b_5[I_5 - 1]^2) - 1]$ | $1, 1, 2, w_{1,2}, w_{2,2}$<br>$5, 2, 2, w_{1,16}, w_{2,16}$ |
| $a_1 = 9.63 \text{ kPa}, b_1 = 1.95$<br>$a_5 = 0.14 \text{ kPa}, b_5 = 1.65$ | $w_{1,2} = 1.95, w_{2,2} = 2.47 \text{ kPa}$<br>$w_{1,16} = 1.65, w_{2,16} = 0.04 \text{ kPa}$ |
| **node 6 - exponential linear second invariant**<br>**node 16 - exponential quadratic fifth invariant** |  |
| $\psi = \frac{1}{2}a_2/b_2[\exp(b_2[I_2 - 3]) - 1]$<br>$+ \frac{1}{2}a_5/b_5[\exp(b_5[I_5 - 1]^2) - 1]$ | $2, 1, 2, w_{1,6}, w_{2,6}$<br>$5, 2, 2, w_{1,15}, w_{2,16}$ |
| $a_2 = 8.30 \text{ kPa}, b_2 = 1.15$<br>$a_5 = 0.14 \text{ kPa}, b_5 = 1.64$ | $w_{1,6} = 1.15, w_{2,6} = 3.62 \text{ kPa}$<br>$w_{1,16} = 1.64, w_{2,16} = 0.04 \text{ kPa}$ |
| **node 6 - exponential linear second invariant**<br>**node 14 - exponential linear fifth invariant** |  |
| $\psi = \frac{1}{2}a_2/b_2[\exp(b_2[I_2 - 3]) - 1]$<br>$+ \frac{1}{2}a_5/b_5[\exp(b_5[I_5 - 1]) - 1]$ | $2, 1, 2, w_{1,6}, w_{2,6}$<br>$5, 1, 2, w_{1,14}, w_{2,14}$ |
| $a_2 = 8.21 \text{ kPa}, b_2 = 1.16$<br>$a_5 = 0.04 \text{ kPa}, b_5 = 3.49$ | $w_{1,2} = 1.16, w_{2,2} = 3.54 \text{ kPa}$<br>$w_{1,14} = 3.49, w_{2,14} = 0.01 \text{ kPa}$ |
| **node 2 - exponential linear first invariant**<br>**node 14 - exponential linear fifth invariant** |  |
| $\psi = \frac{1}{2}a_1/b_1[\exp(b_1[I_1 - 3]) - 1]$<br>$+ \frac{1}{2}a_5/b_5[\exp(b_5[I_5 - 1]) - 1]$ | $1, 1, 2, w_{1,2}, w_{2,2}$<br>$5, 1, 2, w_{1,14}, w_{2,14}$ |
| $a_1 = 8.62 \text{ kPa}, b_1 = 2.38$<br>$a_5 = 0.13 \text{ kPa}, b_5 = 2.21$ | $w_{1,6} = 2.38, w_{2,6} = 1.81 \text{ kPa}$<br>$w_{1,14} = 2.21, w_{2,14} = 0.03 \text{ kPa}$ |
| **node 1 - linear first invariant - Holzapfel**<br>**node 12 - exponential quadratic fourth invariant** |  |
| $\psi = \frac{1}{2}\mu[I_1 - 3]$<br>$+ \frac{1}{2}a/b[\exp(b[I_4 - 3]^2) - 1]$ | $1, 1, 1, w_{1,1}, w_{2,1}$<br>$4, 2, 2, w_{1,12}, w_{2,12}$ |
| $\mu = 12.90 \text{ kPa}$<br>$a = 1.98 \text{ kPa}, b = 6.59$ | $w_{1,1} = 6.45, w_{2,1} = 1.00 \text{ kPa}$<br>$w_{1,12} = 6.59, w_{2,12} = 0.15 \text{ kPa}$ |

**Figure 5:**
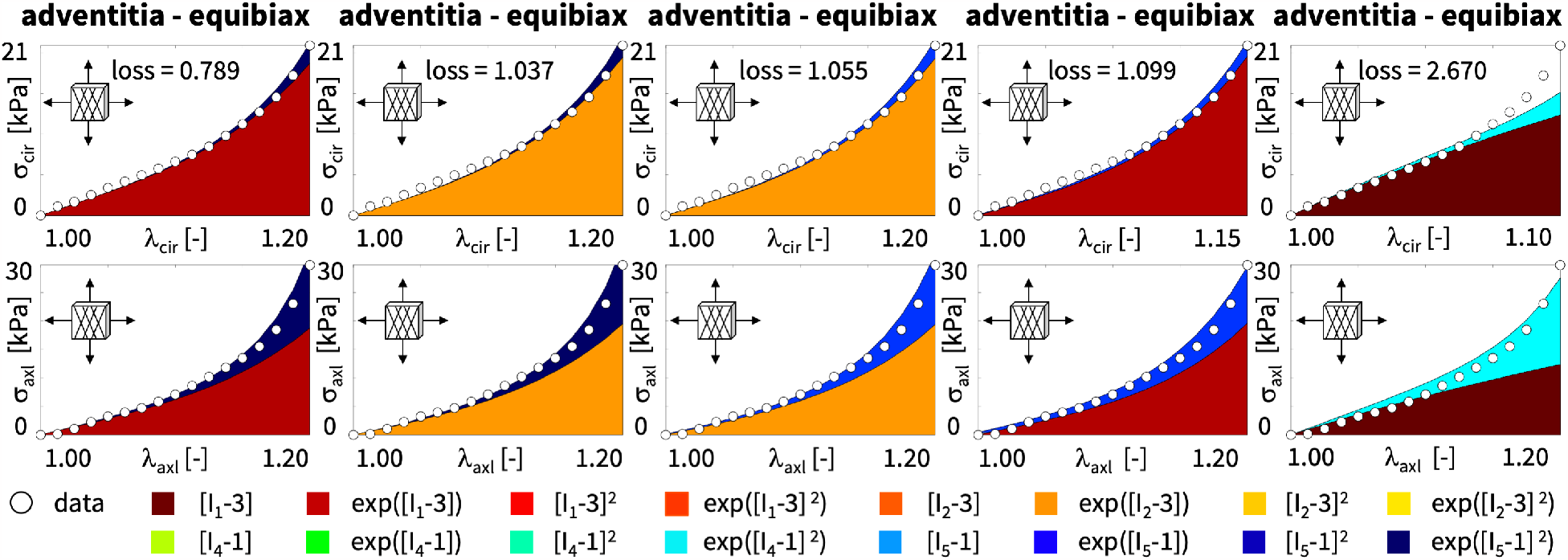
Best-in-class two-term models of the adventitia. True stresses *σ*_cir_ and *σ*_axl_ as functions of stretches *λ*_cir_ and *λ*_axl_ of the constitutive neural network from Figure 2, trained with all ten datasets from Table 2 simultaneously. The first four columns illustrate the best-in-class two-term models from Figure 2, right; the right column illustrates the Holzapfel model [21] for comparison. Circles represent the equibiaxial testing data from Table 2. Color-coded regions represent the discovered model terms. The remaining loss indicates the quality of the overall fit.

### Discovering the best model and parameters

Next, we discover the best model and parameters–but now without prescribing the number of terms–and use the model to validate simulations with our universal material subroutine against the experimental data and against the stress plots from our model discovery. Model discovery is a sophisticated trade-off between the number of discovered terms and the accuracy of the fit [38]. Fortunately, we can fine-tune this trade-off by adding an *L*_*p*_ regularization term to the loss function in equation (14). Specifically, with *L*_1_ regularization and a penalty parameter *α* varying between *α* = [0.000, 0.001, 0.010, 0.100], we observe that we can tune the number of discovered model terms between five and one. For our example, a penalty parameter of *α* = 0.001 provides a good balance between the number of terms and the accuracy of the fit. Strikingly, for this penalty parameter, the network discovers *exactly the same model* for the media and the adventitia: a three-term model with the isotropic linear and exponential first invariant terms and the anisotropic quadratic fifth invariant term,

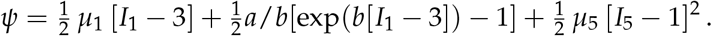

While the discovered *model is the same* for both tissue types, the discovered *parameters are different*, with *μ*_1_ = 33.45 kPa, *a* = 3.74 kPa, *b* = 6.66, *μ*_5_ = 2.17 kPa for the media and *μ*_1_ = 8.30 kPa, *a* = 1.42 kPa, *b* = 6.34, *μ*_5_ = 0.49 kPa for the adventitia. These parameters reflects the different tissue compositions [23], with the media about three to four times stiffer than the adventitia. The discovered model translates into the following four-line parameter table for our universal material subroutine,

~~~
   *PARAMETER TABLE,TYPE=“UNIVERSAL_TAB”
   1,1,1,w_1,1_,w_2,1_ 1,1,2,w_1,2_,w_2,2_ 5,2,1,w_1,15_,w_2,15_ 9,2,1,w_1,15_,w_2,15_
~~~

with `w`_1,1_ = 38.01, `w`_2,1_ = 0.44 kPa, `w`_1,2_ = 6.66, `w`_2,2_ = 0.28 kPa, `w`_1,15_ = 24.68, `w`_2,15_ = 0.04 kPa for the media and `w`_1,1_ = 34.28, `w`_2,1_ = 0.12 kPa, `w`_1,2_ = 6.34, `w`_2,2_ = 0.11 kPa, `w`_1,15_ = 15.32, `w`_2,15_ = 0.02 kPa for the adventitia.

Figures 6 and 7 illustrate the discovered model for the media and the adventita, top, and, for validation, the finite element simulations with our universal material subroutine, bottom. The circles illustrate the biaxial testing data from Tables 1 and 2. The color coded regions highlight the contributions of the individual model terms to the circumferential and axial stresses, *σ*_cir_ and *σ*_axl_, as functions of stretches, *λ*_cir_ and *λ*_axl_. The dark red and red regions represent the isotropic linear first invariant neo Hooke term [59] and the exponential first invariant Demiray term [9]. The blue regions represent the anisotropic quadratic fifth invariant term. Overall, the discovered model provides an excellent fit to the data, both for the media and the adventitia. In both examples, in Figures 6 and 7, the finite element simulations with our universal material subroutine, bottom, agree well with the experimental data and with the model discovery plots, top.

**Figure 6:**
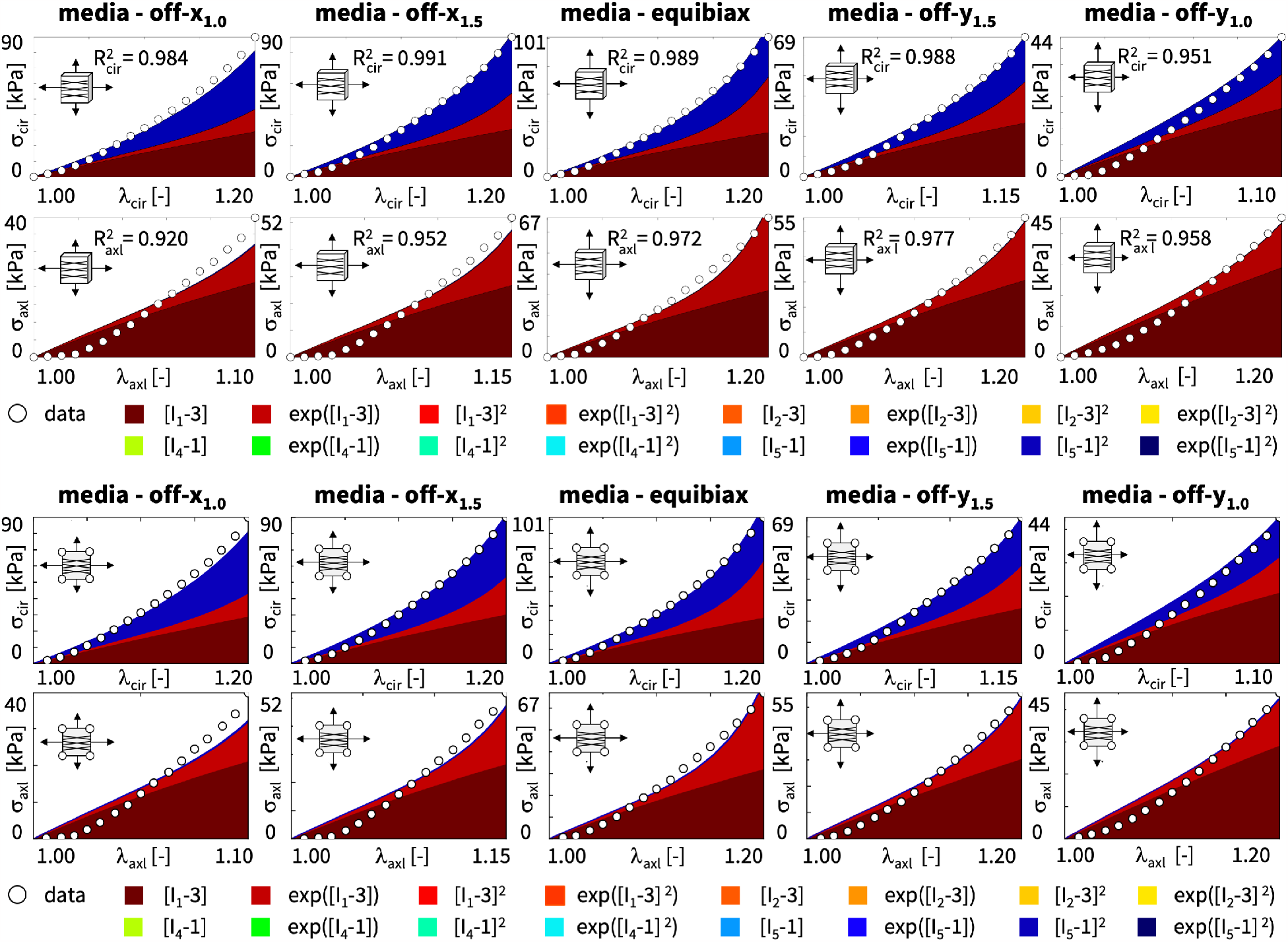
Discovered model and finite element simulation of the media. True stresses *σ*_cir_ and *σ*_axl_ as functions of stretches *λ*_cir_ and *λ*_axl_ of the constitutive neural network from Figure 2, trained with all ten datasets from Table 1 simultaneously. Circles illustrate the biaxial testing data from Table 1. Top graphs display the discovered model, 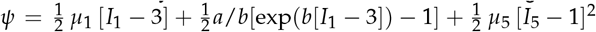 bottom graphs display the finite element simulation with the discovered parameters *μ*_1_ = 33.45 kPa, *a* = 3.74 kPa, *b* = 6.66, *μ*_5_ = 2.17 kPa.

**Figure 7:**
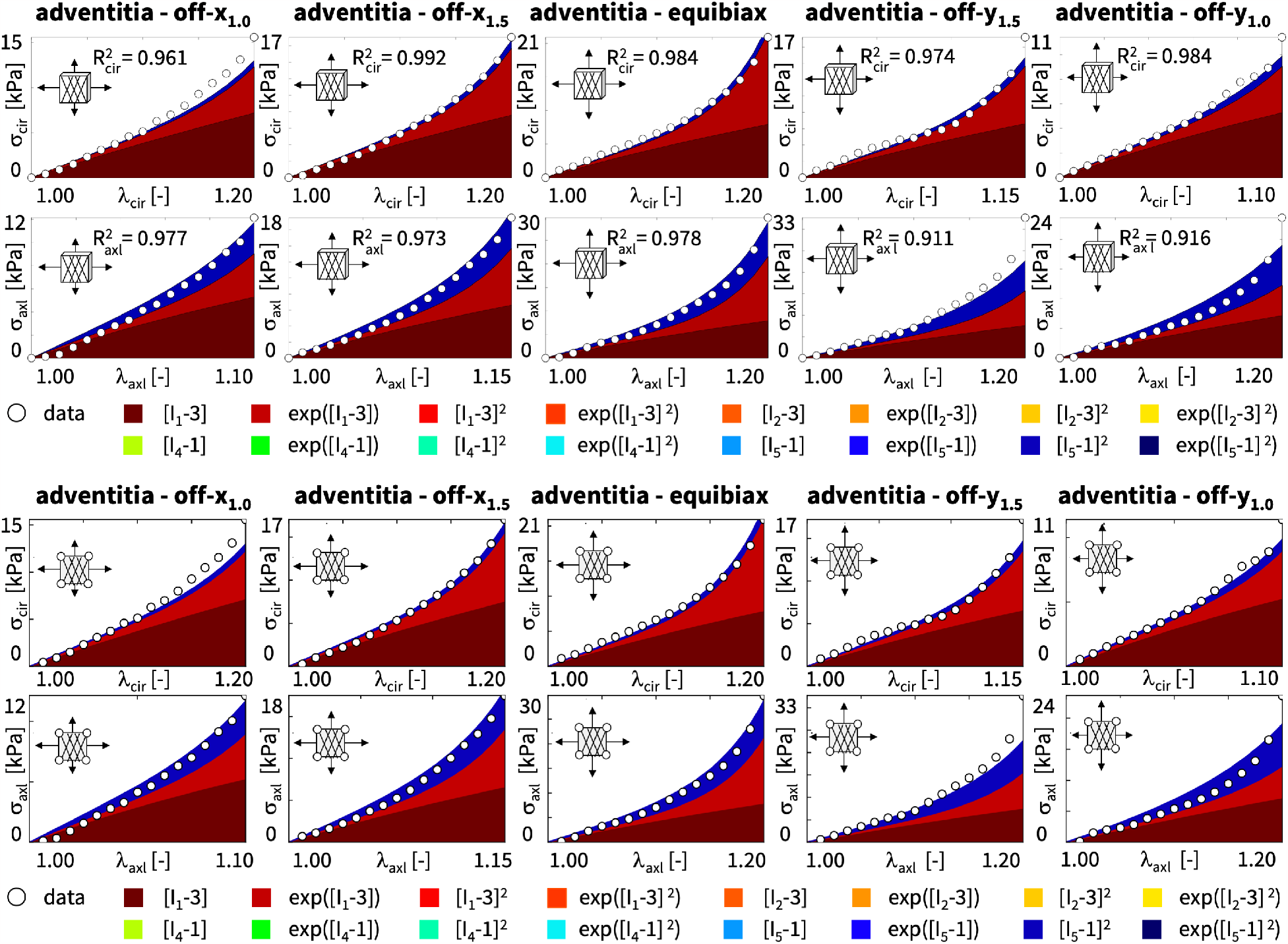
Discovered model and finite element simulation of the adventitia. True stresses *σ*_cir_ and *σ*_axl_ as functions of stretches *λ*_cir_ and *λ*_axl_ of the constitutive neural network from Figure 2, trained with all ten datasets from Table 2 simultaneously. Circles illustrate the biaxial testing data from Table 2. Top graphs display the discovered model, 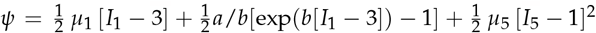 bottom graphs display the finite element simulation with the discovered parameters *μ*_1_ = 8.30 kPa, *a* = 1.42 kPa, *b* = 6.34, *μ*_5_ = 0.49 kPa.

### Predicting wall stresses in the human aortic arch

To explore whether our finite element simulations generalize robustly, from the material point level to the structural level, we now use our universal material subroutine to predict the wall stresses across the human aortic arch and compare our results against the Holzapfel model [17]. We explore the aortic arch during diastole, at a blood pressure of 80 mmHg, and during systole, at 120 mmHg, and visualize the predicted stresses and stretches in the media, in the adventitia, and in selected cross sections. Figure 8 shows our finite element model of the aortic arch, created from high-resolution magnetic resonance images of a healthy, 50th percentile U.S. male [43, 44]. We assume an average aortic wall thickness of 3.0 mm, where the inner 75% of the wall make up the media and the outer 25% make up the adventitia. The finite element discretization uses 60,684 linear tetrahedral elements for the media and 30,342 linear tetrahedral elements for the adventitia, and has a total of 61,692 degrees of freedom. The local collagen fiber angles against the circumferential direction are *±*7.00*°* in the media and *±*66.78*°* in the adventitia. The simulation in Figure 8, left, uses our newly discovered three-term model with an isotropic linear first invariant neo Hooke term, an isotropic exponential first invariant Demiray term, and an anisotropic quadratic fifth invariant term, 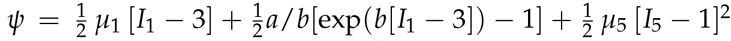 with the discovered parameters *μ*_1_ = 33.45 kPa, *a* = 3.74 kPa, *b* = 6.66, *μ*_5_ = 2.17 kPa for the media and *μ*_1_ = 8.30 kPa, *a* = 1.42 kPa, *b* = 6.34, *μ*_5_ = 0.49 kPa for the adventitia. The simulation in Figure 8, right, uses the Holzapfel model with an isotropic linear first invariant neo Hooke term and an anisotropic exponential term that couples the first and fourth invariants through the dispersion parameter *κ*, 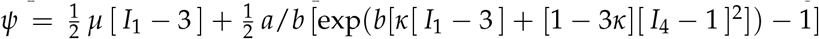 with the best-fit parameters *μ* = 48.68 kPa, *a* = 6.67 kPa, *b* = 23.17, *κ* = 0.074 for the media and *μ* = 13.22 kPa, *a* = 0.93 kPa, *b* = 12.06, *κ* = 0.091 for the adventitia.

**Figure 8:**
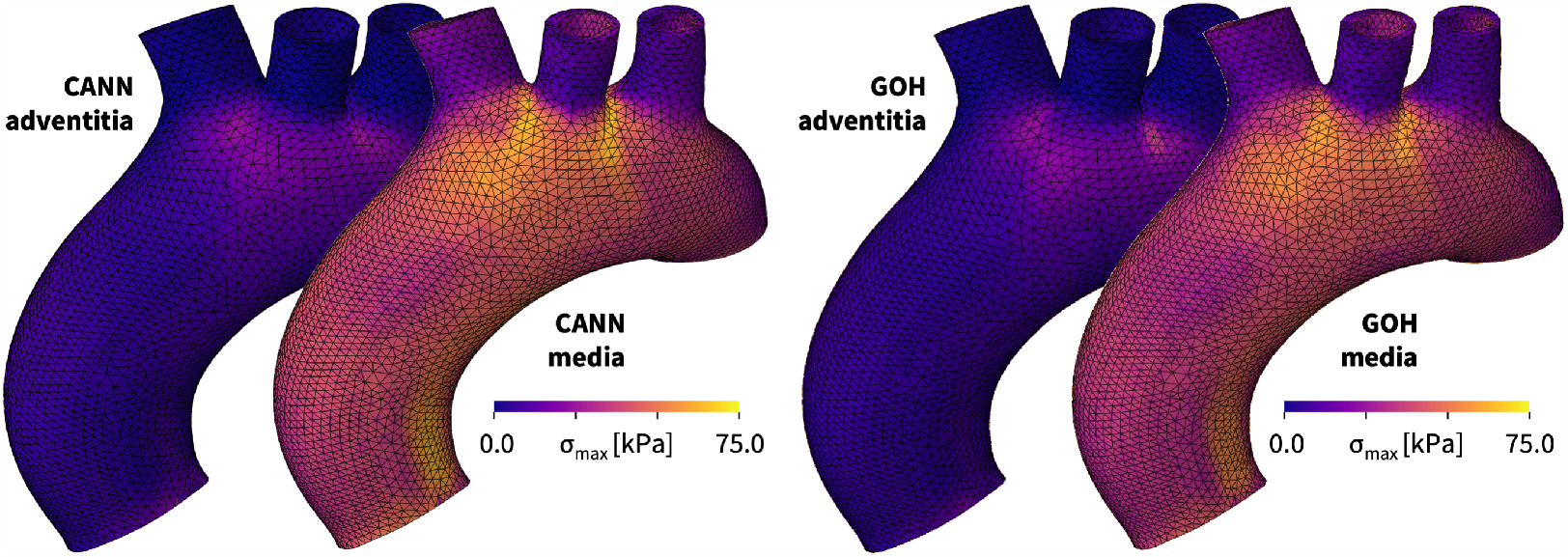
Human aortic arch model and wall stresses in the media and adventitia. The finite element discretization uses linear tetrahedral elements, 60,684 for the media and 30,342 for the adventitia, and has a total of 61,692 degrees of freedom. The color code highlights the maximum principal stresses in the media and adventitia of the human aortic arch predicted by the newly discovered three-term model, left, and by the Holzapfel model, right, both trained with the media and adventitia datasets from Tables 1 and 2.

Figure 8 illustrates four stress profiles that provide a first glance at the performance of both models: First, we emphasize that all large scale structural simulations with our universal material subroutine run and converge robustly, and predict physically reasonable, smooth stress profiles across the entire aortic arch. Second, we observe that, for both constitutive models, the maximum principal stresses in the media are about three times higher than in the adventitia, which agrees well with the recorded stresses in the experiment in Tables 1 and 2 and with the discovered stiffness-like parameters in Tables 3 through 5. Third, and most interestingly, the direct side-by-side comparison of the two different models reveals an excellent agreement of the stress profiles in the low-stress regimes of the adventitia, and a very good agreement in the high-stress regimes of the media, with only a few minor local discrepancies. Overall, we conclude that our universal material subroutine generalizes well from the local material point level to the global structural level and that the simulations with our newly discovered three-term model perform similar to the widely used Holzapfel model [17].

### Predicting aortic arch mechanics with the newly discovered model

Figure 9 illustrates the circumferential and radial stresses and stretches in the media, in the adventita, and in selected cross sections, during diastole, top, and systole, bottom. All simulations use the newly discovered model, 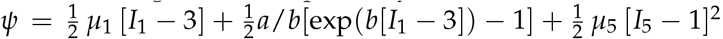 with the discovered parameters *μ*_1_ = 33.45 kPa, *a* = 3.74 kPa, *b* = 6.66, *μ*_5_ = 2.17 kPa for the media and *μ*_1_ = 8.30 kPa, *a* = 1.42 kPa, *b* = 6.34, *μ*_5_ = 0.49 kPa for the adventitia. The simulations provide a nuanced perspective of the mechanics of the aortic arch and detailed insights into the performance of the new three-term model: First, we note that both, stresses and stretches, are larger during systole than during diastole, larger in the the media than in the adventitia, and larger circumferentially than axially. Second, in the stresses profiles, we observe a significant jump between the media and adventita layers, which is most visible in the cross sectional view, and most pronounced during systole. These intra-layer stress discontinuities could play a critical role in the pathogenesis of aortic dissection and aortic aneurysm formation. Third, in the stretch profiles, we observe regional peaks beyond the experimental testing and network training regime of 1.0≤*λ*≤1.2, which are highlighted in bright yellow and most prominent in the circumferential stretch during systole. The smooth stress and stretch profiles beyond the training regime suggest that the discovered model generalizes well to larger stretch regimes, 1.2≤*λ*, and to higher blood pressures. Overall, we conclude that our newly discovered model can predict physically meaningful stretch and stress profiles in complex biological structures and accurately capture the local and global mechanics of the aortic wall.

**Figure 9:**
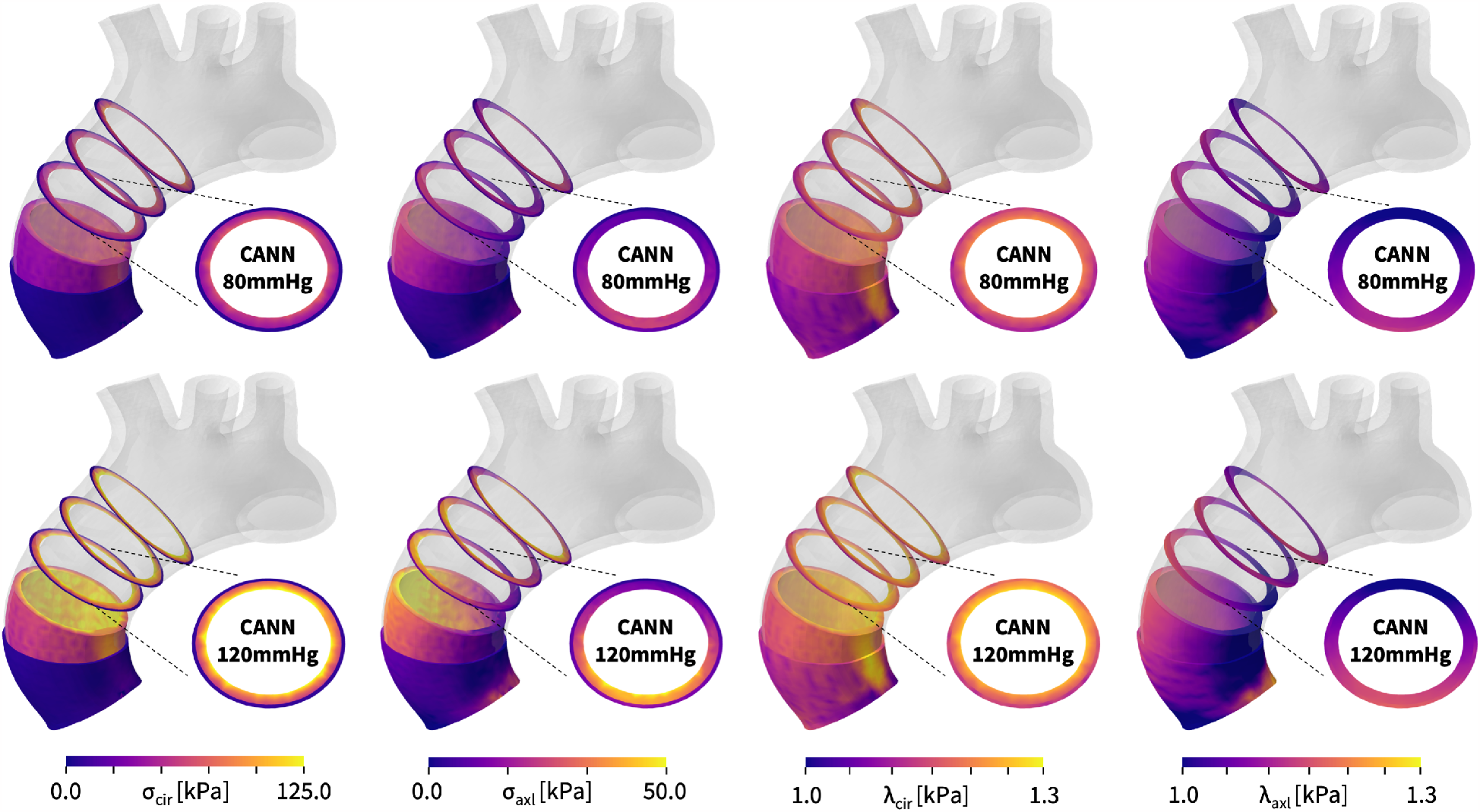
Diastolic and systolic stresses and stretches in the human aortic arch predicted by the newly discovered model. Circumferential and radial stresses, *σ*_cir_ and *σ*_axl_, and stretches, *λ*_cir_ and *λ*_axl_, in the media, in the adventita, and in selected cross sections, during diastole, top, and systole, bottom. Simulations use the discovered model, 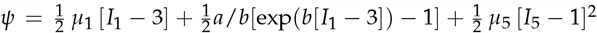 with the discovered parameters *μ*_1_ = 33.45 kPa, *a* = 3.74 kPa, *b* = 6.66, *μ*_5_ = 2.17 kPa for the media and *μ*_1_ = 8.30 kPa, *a* = 1.42 kPa, *b* = 6.34, *μ*_5_ = 0.49 kPa for the adventitia.

### Predicting aortic arch mechanics with the Holzapfel model

Figure 10 illustrates the stresses and stretches in the in the media, in the adventita, and in selected cross sections, during diastole and systole similar to Figure 9, but now using the Holzapfel model [17], 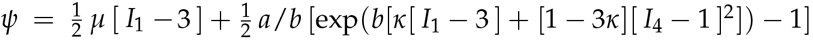with the best-fit parameters, *μ* = 48.68 kPa, *a* = 6.67 kPa, *b* = 23.17, *κ* = 0.074 for the media and *μ* = 13.22 kPa, *a* = 0.93 kPa, *b* = 12.06, *κ* = 0.091 for the adventitia. The simulations provide additional insight into the similarities and differences of both constitutive models: First, within the experimental testing and network training regime, 1.0≤*λ*≤1.2, the predictions with the Holzapfel model in Figure 10 are virtually identical to the predictions with our new three-term model in Figure 9. This is particularly evident during diastole, and across the entire adventita during both diastole and systole. Second, beyond the experimental testing and network training regime, 1.2≤*λ*, we observe small discrepancies between both models, which are located primarily in the bright yellow regions of the high-stretch regime. This agrees with our intuition that the exponential term of the Holzapfel model introduces a more pronounced stiffening than the quadratic term of our newly discovered model, especially in the high-stretch regime. Overall, we conclude that both models perform almost identically during diastole, within their training regime, and very similarly during systole, beyond their training regime, where the stresses of the Holzapfel model are locally slightly higher than those of the new three-term model, while its stretches are locally slightly lower.

**Figure 10:**
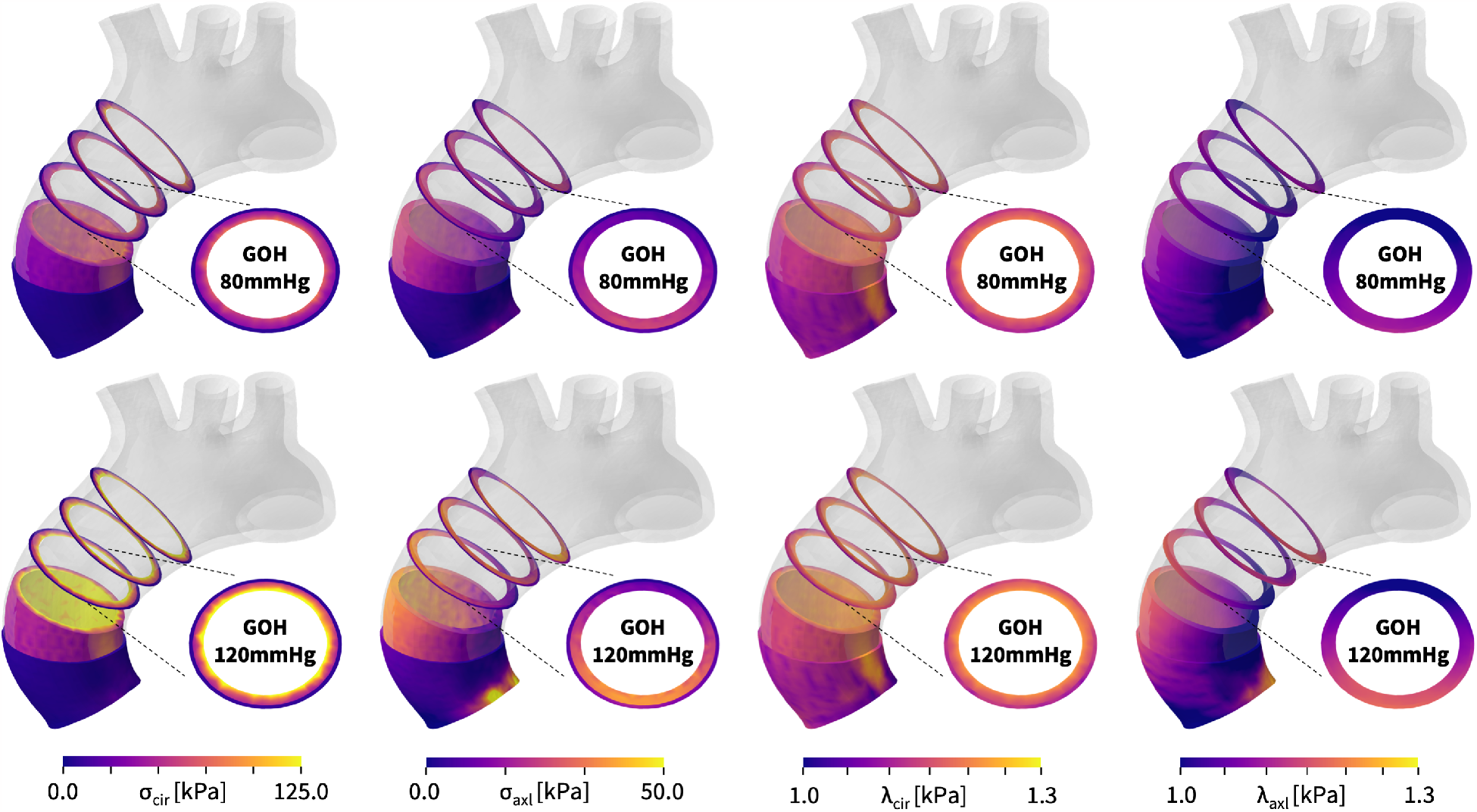
Diastolic and systolic stresses and stretches in the human aortic arch predicted by the Holzapfel model. Circumferential and radial stresses, *σ*_cir_ and *σ*_axl_, and stretches, *λ*_cir_ and *λ*_axl_, in the media, in the adventita, and in selected cross sections, during diastole, top, and systole, bottom. Simulations use the Holzapfel model, 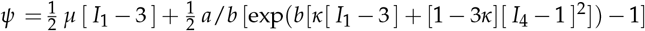 with the best-fit parameters, *μ* = 48.68 kPa, *a* = 6.67 kPa, *b* = 23.17, *κ* = 0.074 for the media and *μ* = 13.22 kPa, *a* = 0.93 kPa, *b* = 12.06, *κ* = 0.091 for the adventitia.

## 5 Discussion

Computational modeling is vital for unraveling the biomechanics of the aorta and offering insights into its disease mechanisms. Finite element analyses enable a precise identification of regions of non-physiological deformations or stresses, which could indicate the onset of vascular diseases such as aortic dissection or aneurysm formation. Constitutive modeling lies at the heart of any finite element analysis, and selecting the best model and parameters is crucial for its success. Common finite element analysis tools offer a wide variety of constitutive models to choose from, but selecting the best model remains a matter of user experience and personal preference. *The objective of this study is to eliminate user bias and fully automate the process of model selection using constitutive neural networks*. We train these networks with experimental data from biaxial extension tests on the human aortic media and adventita, and discover the best models and parameters to explain the data. Our model discovery workflow automatically generates an input file for a universal material subroutine that can represent more than 60,000 different constitutive models and is seamlessly embedded in the finite element analysis pipeline. In this manuscript, we rationalize the process of model discovery, discuss models of different complexity, demonstrate their performance on the experimental data, compare them against the current gold standard model, and use the best model and parameters to predict stress and stretch profiles across the aortic arch during diastole and systole.

### Model discovery is a balance between complexity and accuracy

The *universal approximation theorem* states that a neural network with a single hidden layer–with a sufficient number of nodes and appropriate activation functions–can approximate *any continuous function* on a compact subset of its domain to arbitrary precision [27]. This implies that, with a sufficient number of nodes, our network should be able to approximate any of the stretch-stress pairs of our biaxial tests. However, in constitutive modeling, we are not interested in just learning *any* function. Instead, we seek to discover the best function that not only approximates the data, but also satisfies common thermodynamic principles and physical constraints [33]. These include material objectivity, material symmetry, incompressibility, polyconvexity [30, 54], and thermodynamic consistency [32, 56]. Conveniently, we hardwire these principles into our constitutive neural network in Figure 2 to ensure that our discovered functions satisfy these constraints a priori. Specifically, our network has two hidden layers and represents the free energy function as the sum of the contributions of the sixteen nodes of its second layer [35]. Naturally, activating all sixteen nodes is the best strategy to fine-tune the fit to the data and achieve the highest level of accuracy. At the same time, the resulting sixteen-term model is inherently complex and difficult to interpret [52]. Nonetheless, if we are *only* interested in finding the *best-fit model* and parameters for a finite element analysis, this is probably just fine. We can feed all sixteen terms directly into our universal material subroutine and perform our engineering analysis. Undoubtedly, this will make the best and most explicit use of the available data.

### Model discovery can be non-robust and non-interpretable

In many practical applications, we are not just interested in finding the best-fit model with an arbitrarily large number of terms. Instead, we want to *discover the most relevant terms* to best describe experimental data. This can have multiple reasons: First, minimizing the loss function (14) with 16 terms and 24 independent weights translates into a complex *non-convex optimization* problem with flat gradients and multiple local minima [38]. It is computationally expensive, if not impossible, to find its global minimum. Second, with so many degrees of freedom, there is a *risk of overfitting*. Even if we found the global minimum, it might be highly *sensitive to outliers* or measurement errors [6]. In other words, we might find the best-fit model for a specific data set, but this model tends to be *non-robust* and *non-generalizable* to unseen data. Third, and probably most importantly for our purposes, a sixteen-term model is virtually *non-interpretable* [11]. We cannot interpret the relevance of its terms, compare the meaning of its parameters, and identify the underlying mechanisms associated with individual terms. A fully activated model provides virtually no microstructural insights into the material response. This raises the holy grail question in model discovery: How can we fine tune the number of terms?

### Lp regularization promotes robust and interpretable models

The concept of *L*_*p*_ regularization or bridge regression dates back more than three decades and was introduced to shrink the parameter space in a data analysis [13]. It has re-gained attention as a powerful tool to promote sparsity in system identification [5], and, most recently, in discovering constitutive models from data [11, 38]. *L*_*p*_ regularization adds a penalty term, 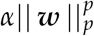, to the loss function (14), where *α* ≥ 0 is a non-negative penalty parameter and 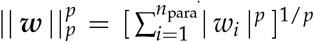 is the *L*_*p*_ norm of the vector of network weights ***w***. *L*_*p*_ regularization introduces two hyperparameters, the power *p* by which it penalizes the individual model parameters, and the penalty parameter *α* by which it scales the relative importance of the regularization term compared to the network loss [38]. Both parameters enable a precise control of model discovery and it is crucial to understand their mathematical subtleties, computational implications, and physical effects: *L*_0_ regularization or *subset selection* directly penalizes the number of non-zero terms by solving the discrete combinatorial problem, which is a simple and unbiased method to explicitly prescribe the number of terms [29]. *L*_1_ regularization or *lasso* enables feature selection and induces sparsity by reducing some weights exactly to zero, which effectively reduces model complexity and improves interpretability [58]. *L*_2_ regularization or *ridge regression* seeks to reduce outliers and improve predictability by reducing absolute values while maintaining all parameters [18], which essentially does the opposite of what we seek to accomplish here.

### L0 regularization identifies the best-in-class models

*L*_0_ regularization or *subset selection* turns the continuous model selection problem into a *discrete combinatorial problem* with (2^*n*^-1) possible combinations of terms [29]. This makes this type of regularization computationally intractable for models with a large number of terms. However, instead of performing a subset selection from all possible 65,535 models, we use it to discover the best-in-class one- and two-term models–out of subsets of 16 and 120 possible models–and gain insight into the relevant terms and model parameters [38]. Interestingly, in our example, for both media and adventitia, the best-in-class one-term model in Table 3, with the lowest remaining loss of 84.49 for the media and 3.29 for the adventita, is the classical exponential linear first invariant Demiray model [9]. The best-in-class two-term models in Table 4 for the media and Table 5 for the adventitia expand this term by either the quadratic fifth invariant or the exponential quadratic fifth invariant, with remaining losses of 8.84 for the media and 0.79 for the adventitia. Since *L*_0_ regularization explicitly penalizes every additional term in the loss function (14) by *α*, it favors the two-term model for the media for *α*≤84.49/8.84 = 9.56 and for the adventitia for *α*≤3.29/0.79 = 4.17, and only selects the purely isotropic one-term Demiray model [9] for penalty parameters larger than these values. This simple example illustrates the role of the penalty parameter *α* as a hyperparamter to fine-tune subset selection by modulating the number of non-zero terms.

### L1 regularization induces sparsity and improves interpretability

A less invasive approach to regularize the loss function without having to explicitly probe all combinations of terms is *L*_1_ regularization or lasso [58]. Here we apply *L*_1_ regularization with varying penalty parameters *α* and monitor the remaining loss. We find a reasonable balance of complexity and accuracy for a penalty parameter of *α* = 0.001. Strikingly, for this parameterization, out of all 65,535 combinations of terms, our network discovers *exactly the same* model for the media and the adventitia: a three-term model with an isotropic linear first invariant neo Hooke term [59], an exponential linear first invariant Demiray term [9], and the anisotropic quadratic fifth invariant term. Its non-zero network weights translate into interpretable material parameters in the form of a shear modulus, a stiffness-like parameter, an exponential coefficient, and a shear-type modulus, all with physically meaningful units. *Our newly discovered model is sparse, robust, and interpretable, its contains terms of popular constitutive models, and is a natural generalization of our discovered best-in-class one- and two-term models*. We feel that it strikes an excellent balance between complexity and accuracy. From Figures 6 and 7, we conclude that it approximates our experimental data well and integrates seamlessly into our universal material subroutine.

### Our discovered model generalizes from the material point level to structural analysis

One of the main reasons to develop constitutive models for biological tissues is to perform realistic biomedical simulations [42, 44]. The ultimate test of our discovered model is to probe its performance in realistic finite simulations, beyond the material point level. Here we use the example of stress analysis in the aortic arch. Understanding the structural and mechanical distinctions between the media and adventitia layers of the aorta is crucial for comprehending vascular health and disease [23, 28]: The media is rich in elastin fibers and smooth muscle cells, it provides elasticity and contractility, and enables hemodynamic function. The adventitia consists primarily of collagen fibers and fibroblasts and provides structural support and integrity. By building our model *directly from data*–without user bias through model selection–we can precisely capture the nuances between the load carrying capacity of the media and the adventitia [40, 41]. Disruptions in the delicate balance between these layers contribute to pathological conditions such as aortic dissection or aneurysms formation [47, 50]. Mechanical heterogeneity and regional stress variations play a pivotal role at the onset of these conditions. Our finite element model is built around our new universal material subroutine [45] that can account for these layer-specific properties and aid in predicting disease progression, assessing rupture risk, and developing targeted interventions. This subroutine not only includes our discovered three-term model, but all 65,535 possible models of our constitutive neural network in Figure 2 [35], simply by twenty-four lines of its input file. A side-by-side comparison with the popular Holzapfel model in Figures 9 and 10 suggests that our discovered model not only performs *identically during diastole*, within the stretch range of the training regime, and but also performs *nearly similarly during systole*, beyond the initial training regime. The small local discrepancies between both models are not a flaw of our new model, but rather a result of the *limited experimental test range* within stretches of only 1.0 to 1.2. From the experimental stretch-stress curves in Figure 1, we conclude that within this range, the stress response of the fibers is neither fully quadratic as in our discovered model, nor fully exponential as in the Holzapfel model [17]. Overall, we believe that newly discovered model performs well in realistic structural simulations and can provide a comprehensive understanding of the interplay between the layers of the aorta to informs strategies for early disease detection, risk stratification, and tailored therapeutic approaches in the benefit of cardiovascular health.

### Limitations

While our results solidly suggest that we can discover interpretable models with physically meaningful parameters from data and integrate these models into a finite element simulation via our new universal material subroutine, a few limitations remain: First, here we have prototyped our approach for discovering a personalized arterial model of a healthy 56-year-old male. We are currently expanding our method to include all *n* = 17 healthy and *n* = 11 aneurysmatic aortas of the initial study [40]. Second, while our study shows that *L*_*p*_ regularization is a robust method to control the number of model terms through the penalty parameter *α*, especially the low-penalty models with a large number of discovered terms remain sensitive to the initial conditions. If the goal is to discover *the best model* with a small number of interpretable terms, we recommend to always perform an *L*_0_ regularization first, and solve for the discrete combinatorics problem–at least for the best-in-class one- and two-term models–to gain a feeling for the relevant terms [38]. If the goal is to discover *a viable model* for a finite element simulation, the sparseness of the solution is less relevant, since we can feed any discovered model into our universal material subroutine and obtain comparable results. Third, while different discovered models perform similarly within the training range, they may deviate outside the training regime. For finite element simulations, this may occur in regions of local stress concentrations, where the simulated stretches and stresses exceed the experimental measurement range. This is not a flaw of the model discovery itself, but rather a limitation of the available training data, which, in our example, did not properly tease out the stretch-stiffening regime. As a result, the discovered anisotropic term that best explains our available data turns out to be quadratic, and not exponential like in the classical Holzapfel model [17, 21]. Fourth, for illustrative purposes, the neural network and the material subroutine we propose here are intentionally invariant-decoupled. We are currently finalizing an advanced material subroutine that can handle coupled invariants like *I*_1_ and *I*_4_ [17], selective activation under tension only [17], and account for quasi-incompressibility through the third invariant *I*_3_. Finally, to address the current limitation to hyperelastic materials, we have recently expanded the concept of constitutive neural networks to viscoelasticity [62] and to general inelasticity [20].

## 6 Conclusion

Personalized computational simulations can help us understand the biomechanics of cardiovascular disease, predict patient-specific disease progression, and personalize treatment and intervention. Material modeling is critical to realistic physics-based simulations, but selecting the best model is limited to a few highly trained specialists in the field. In biomedical applications, poor model selection does not only jeopardize the success of the entire simulation, but can have life-threatening consequences for the patient. Here we explore the feasibility of removing user involvement and automating material modeling in finite element analyses. We leverage recent developments in constitutive neural networks, machine learning, and artificial intelligence to discover the best constitutive model from thousands of possible combinations of a few functional building blocks. We seamlessly integrate all discoverable models into the finite element workflow by creating a universal material subroutine that contains more than 60,000 models, made up of 16 individual terms. Our results suggest that constitutive neural networks can robustly discover various flavors of arterial models from data, feed these models directly into a finite element simulation, and predict stress and strain profiles that compare favorably to the classical Holzapfel model. Replacing dozens of individual material subroutines by a single universal material subroutine will make finite element simulations more accessible and user-friendly, more robust and reliable, and less vulnerable to human error. Democratizing biomedical simulation by automating model selection could induce a paradigm shift in physics-based simulation, broaden access to simulation technologies, and empower individuals with varying levels of expertise and diverse backgrounds to actively participate in scientific discovery in the benefit of human health.

## >Acknowledgements

We acknowledge support through the NSF CMMI Award 2320933 Automated Model Discovery for Soft Matter to EK and the NWO Veni Talent Award 20058 to MP.

## Notes

### Competing Interest Statement

The authors have declared no competing interest.

